# Tissue composition shapes differential skeletal integration strategies during axolotl limb regeneration

**DOI:** 10.64898/2026.02.26.708175

**Authors:** Rita Aires, Sean D. Keeley, Kerstin Brandt, Mário Carreira, Doğa Berşan Güneş, Yağız Savcı, Pedro Capucho, Ulrike Anne Friedrich, Andreas Dahl, Can Aztekin, Tatiana Sandoval-Guzmán

**Affiliations:** Department of Internal Medicine III, Center for Healthy Aging, University Hospital and Faculty of Medicine Carl Gustav Carus, TUD Dresden University of Technology, 01307 Dresden, Germany; Center for Regenerative Therapies Dresden (CRTD), Center for Molecular and Cellular Bioengineering (CMCB), TUD Dresden University of Technology, 01307 Dresden, Germany; Paul Langerhans Institute Dresden of the Helmholtz Center Munich, University Hospital and Faculty of Medicine Carl Gustav Carus, TUD Dresden University of Technology, 01307 Dresden, Germany; Abel Salazar Institute of Biomedical Sciences, University of Porto, 4200-465 Porto, Portugal; Faculty of Engineering, University of Porto, 4200-465 Porto, Portugal; Graduate School of Science and Engineering, Yıldız Technical University, 34220 Esenler/Istanbul, Türkiye; NOVA Medical School, Faculdade de Ciências Médicas, NMS, FCM, Universidade NOVA de Lisboa, Campo Mártires da Pátria, 130, 1169-056 Lisboa, Portugal; DRESDEN-concept Genome Center (DcGC), Center for Molecular and Cellular Bioengineering (CMCB) Technology Platform, TUD Dresden University of Technology, 01062 Dresden, Germany; German Center for Diabetes Research (DZD e.V.), 85764 Neuherberg, Germany; Friedrich Miescher Laboratory of the Max Planck Society, 72076 Tübingen, Germany

**Keywords:** Axolotl, Regeneration, Tissue Integration, Osteoclasts, Positional amputations, Chemokine evolution

## Abstract

Limb regeneration requires rebuilding the missing structures and integrating them with stump tissues. Osteoclast-mediated tissue resorption is essential for skeletal integration during regeneration. However, it is unknown if skeletal tissue composition impacts its resorption and, if so, how it is regulated.

Here, we show that osteoclast-mediated skeletal resorption is activated in amputations damaging the diaphysis, but not the epiphysis, of skeletal elements. We found that amputations in the calcified diaphysis trigger the sustained expression of *RANKL* and the chemokine *Loc483,* and that the latter is sufficient to induce osteoclast presence in non-resorbing amputations. Finally, our data suggests that the transcriptomic profile of the apical ectodermal cap is impacted by the tissues injured by the amputation.

Overall, our work reveals that tissue composition at the amputation plane shapes adaptations of the regenerative program, which ultimately contribute to the near-seamless tissue integration of the regenerating axolotl limb regardless of the amputation position.

## Introduction

Limb regeneration in the axolotl (*Ambystoma mexicanum*) is an important model for the study of complex structure re-formation and occurs in sequential, yet largely overlapping phases^1,2^. After an amputation, the wound quickly heals through the establishment of a wound epithelium (WE). The immune response is then established by the migration of immune cells into the stump, where they release important factors and help clear pathogens and cell debris. Meanwhile, cells in the WE proliferate, forming the multilayered apical epithelial cap (AEC)^3^. Together, immune cells and the AEC contribute to tissue histolysis by secreting proteolytic enzymes that extensively remodel the stump tissues^3,4^. Finally, a blastema forms under the AEC, which will then proliferate and ultimately differentiate to regrow the missing tissues of the limb.

Although these processes are well studied, the mechanisms by which newly formed tissues integrate with stump structures remain mostly elusive. Remarkably, tissue integration occurs irrespective of the amputation position in the axolotl limb^2,5,6^, which means that regenerative integration processes are equally effective regardless of the specific tissue composition affected by the injury. This poses an interesting challenge especially for the regenerating skeleton, as a typical limb skeletal element comprises a variety of cell types arranged in distinct conformations along its length, resulting in positional differences in thickness, stiffness, and relative proportions^7^. Likewise, cell and tissue heterogeneity in the skeletal element can greatly differ in the span of a few micrometers, as is evidenced by the diaphyseal (i.e., center) and flanking epiphyseal (i.e., proximal and distal) regions. Epiphyses in the axolotl limb are mostly comprised of chondrocytes, which contribute to the growth of the limb and remain cartilaginous throughout the life of the animal^7^. In contrast, the diaphysis is where the primary ossification center will first develop, which will ultimately be comprised of a mix of hypertrophic chondrocytes, osteoblasts, and osteocytes, as well as periskeletal cells and a surrounding calcified extracellular matrix (ECM)^7^. Moreover, skeletal integration often requires the amalgamation of nascent cartilage with the calcified remains of the amputated skeletal element of the stump^6^. Yet, despite these obvious differences, it remains unclear whether or not the mechanisms of skeletal integration differ according to the tissue composition at the amputation site.

Recently, we demonstrated that osteoclast-mediated tissue resorption is important for integrating the regenerated radius and ulna with the previously existing skeletal elements^6^. Osteoclasts are large, multinucleated cells that have an immune origin^8^. In mammals, these cells differentiate in a stepwise manner from common myeloid progenitors (CMPs)^9,10^, which also generate both macrophages and dendritic cell progenitors^8^. Osteoclasts resorb skeletal tissue by adhering to the bone surface and degrading the calcified matrix by secreting protons (H^+^) and proteolytic enzymes such as Cathepsin K (Ctsk) and Matrix Metalloproteinases (MMPs)^9^. During axolotl limb regeneration, osteoclast-mediated tissue resorption occurs in a short and distinct window of time^6^, which contrasts with the long-lasting resorption observed in mammalian bone fractures^11^. This suggests that unique underlying regulatory mechanisms might be employed in regenerative integration.

In this work, we investigated how osteoclast-mediated skeletal resorption is regulated to promote skeletal integration during limb regeneration. We found that this process is dependent on the composition of the injured tissue, with resorption being triggered specifically by amputations through the calcified diaphysis and primarily used for its regenerative integration. Using a combination of spatial transcriptomics and bulk RNA-seq, we show that the RANK/RANKL system likely orchestrates osteoclast differentiation after diaphysis amputations as early as 3 days post-amputation (dpa). Moreover, we also discovered that the previously unknown chemokine *Loc483* has a role in promoting osteoclast differentiation and/or recruitment. Finally, we observed that the amputation site can induce significant transcriptomic differences in the AEC.

Altogether, by exploring two amputation planes affecting different tissue types, our work shows that tissue composition at the injury site induces adaptations in the histolysis process, the immune response, and in the AEC, which may synergistically promote skeletal integration. This demonstrates that early regenerative mechanisms are tailored to the types of tissues affected by the amputation, resulting in the use of differential skeletal integration strategies during regeneration. Ultimately, such adaptations of the regenerative program make possible not only the successful re-formation of all missing limb tissues, but also their integration with the mature structure regardless of the amputation position within the limb.

## Results

### Skeletal resorption is specific to diaphysis amputations

To explore the mechanisms controlling tissue resorption, we chose two amputation planes that, while impacting the same skeletal elements, affected regions with different cellular and ECM compositions (Fig. 1a, vertical dashed lines; Supplementary Fig. 1a). For this, we used axolotls in which the mineralization of diaphyseal regions was already well-developed, but before it showed the first signs of ossification^7^ (Fig. 1a; Supplementary Fig. 1a, b; Supplementary Data 1). In these, the diaphyses of both radii and ulnas were composed of a robust calcified ECM (Fig. 1a, black arrowhead, in yellow between dashed lines) and periskeletal cells (Fig. 1a, white arrowhead) surrounding a hypertrophic zone containing hypertrophic chondrocytes (Fig. 1a, black arrow). In contrast, their epiphyses were mostly constituted by chondrocytes (Fig. 1a, white arrowhead).

**Fig. 1.**
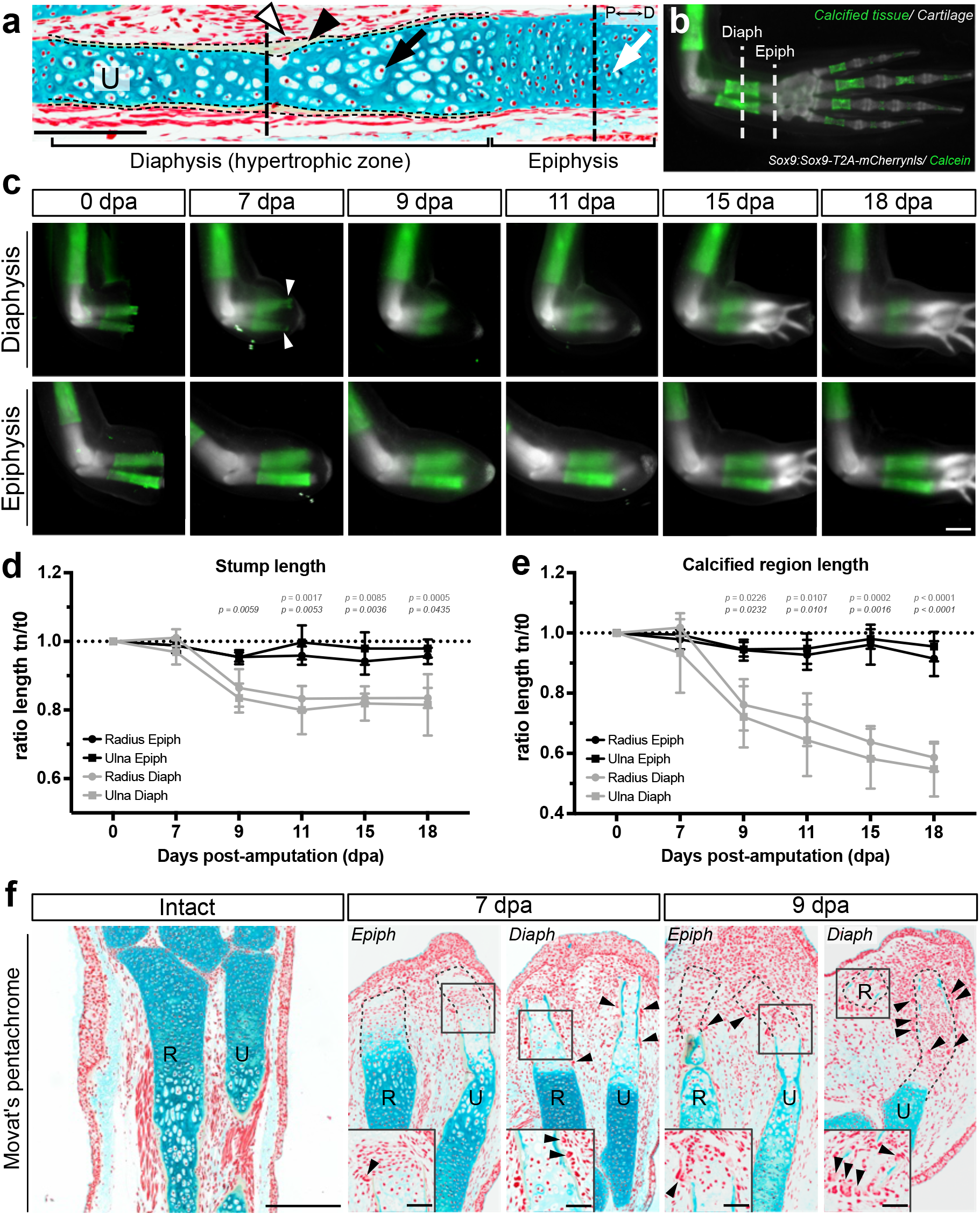
Extensive tissue resorption is specifically triggered by amputations through the calcified diaphyseal region of the radius and ulna. **a** Histological characterization of a longitudinal section of an intact ulna (U) by Movat’s pentachrome. Diaphysis regions comprise calcified ECM (black arrowhead, in yellow between dashed lines), periskeletal cells (white arrowhead) and hypertrophic chondrocytes in the hypertrophic zone (in blue, black arrow). Epiphyseal regions are mostly composed of chondrocytes (in blue, white arrow). Vertical dashed lines represent the approximate sites of amputation planes. Proximal (P) is to the left, while distal (D) is to the right of the image. Representative image of an experiment with n = 3 animals. Scale bar: 300 µm. **b** Schematic representation of amputation planes through the calcified diaphysis and cartilaginous epiphysis in *Sox9-mCherry* animals (labelling chondrocytes, in white) stained with calcein (calcified ECM, in green). **c** Time course of tissue resorption during lower arm regeneration in diaphysis and epiphysis amputations at 0, 7, 9, 11, 15, and 18 dpa. Representative images of an experiment with n = 5 animals per condition. Scale bar: 500 µm. **d** Quantification of the stump length of the radius and ulna after diaphysis and epiphysis amputations over time. **e** Quantification of the calcified region length in radii and ulnas after diaphysis and epiphysis amputations over time. **f** Movat’s pentachrome staining of representative longitudinal sections of intact lower arms and diaphysis- and epiphysis-amputated limbs at 7 and 9 dpa. R, radius, U, ulna. Dashed lines represent the contours of the radius and ulna. Boxes represent the inset location. Black arrowheads indicate multinucleated osteoclasts. Representative image of an experiment with n = 3 animals per time point and amputation plane. Scale bar: 500 µm, scale bar in insets: 100 µm. For c, d, and e, n = 5 animals. For d and e, the lines show mean values over time ± sd. *p*-values in top row refer to diaphysis vs epiphysis amputations in the radius; *p*-values in bottom row refer to diaphysis vs epiphysis amputations in the ulna (two-way ANOVA with Tukey’s post-hoc test). Source data for d and e are provided as a Source Data file.

To accurately assess skeletal tissue resorption, we utilized *Sox9:Sox9-T2a-mCherrynls* (*Sox9-mCherry*) transgenic animals stained with calcein. This allowed for the clear identification of both the cartilaginous skeleton, which expressed *mCherry* in chondrocytes, as well as the calcified diaphysis, which was marked by the binding of calcein to its calcified ECM (Fig. 1b). Using this strategy, we were able to simultaneously assess 1) resorption of the remaining stump skeleton (the length of *mCherry* signal from the elbow joint until the amputation plane); and 2) resorption specifically of the calcified tissue (the length of the calcein-positive region) in the same animals.

Forelimbs of *Sox9-mCherry*/calcein animals were amputated and followed for 18 days. In diaphysis amputated limbs, resorption of the calcified region was first observed at 7 dpa in the form of gaps in the calcein staining (Fig. 1c, top row, white arrowheads). By 9 dpa, the calcified matrix had been significantly resorbed, after which this process considerably slowed up until 18 dpa (Fig. 1c). In contrast, in epiphysis amputations (Fig. 1c, bottom row), both stump skeletal elements and their calcified regions remained relatively unchanged. Quantification of the remaining stump skeleton and respective calcified region length in diaphysis amputations revealed that, on average, approximately 20% of the total length of the stump and up to 40% of the calcified region of radii and ulnas were resorbed by 18 days (Fig. 1d, e). Furthermore, most of this resorption occurred between 7 and 9 dpa, which agreed with previous reports^6^.

Epiphysis amputations, on the other hand, did not exhibit such extensive resorption (Fig. 1c). However, some variations in the length of the stump cartilage and calcified regions at 9 and 11 dpa could be observed (Fig. 1d, e). To further visualize the cellular and ECM structure of tissues after diaphysis and epiphysis amputations, we analyzed the histology of limb sections at 7 and 9 dpa. This showed that, in amputated epiphyses, the cartilaginous matrix of the radius and ulna had undergone considerable remodeling, and that a few of the most distal chondrocytes had even disappeared entirely (Fig. 1f). This explained the observed variations in the length of whole skeletal elements, and matched our previous observations of cartilage undergoing histolysis^12^.

Given their roles as the main cellular effectors of skeletal tissue resorption, we analyzed osteoclast prevalence in the two types of amputations. Diaphysis amputations indeed showed robust osteoclast presence surrounding the calcified region of both skeletal elements at 7 and 9 dpa. In contrast, only a few osteoclasts were observed associated to the skeletal elements in epiphysis amputations at these time points (Fig. 1f, black arrowheads; Fig. 1f, insets). Next, we decided to follow osteoclasts in vivo over time after diaphysis and epiphysis amputations. For that, we combined the reporter line *Sox9-mCherry* with *Ctsk-eGFP* transgenic animals, in which eGFP is driven by the promoter of the mature osteoclast marker *Ctsk*^6^, to generate *Sox9::Ctsk* animals. In vivo imaging confirmed the extensive presence of osteoclasts at 7 and 9 dpa in diaphysis amputations, which quickly decreased at 11 dpa, and was cleared by 15 dpa (Fig. 2a, top row). In contrast, minimal to no osteoclast presence was found in epiphysis amputations (Fig. 2a, bottom row). As expected, we saw no faulty integration phenotypes in either epiphysis- or diaphysis-amputated limbs at 45 dpa (Supplementary Fig. 1c).

**Fig. 2.**
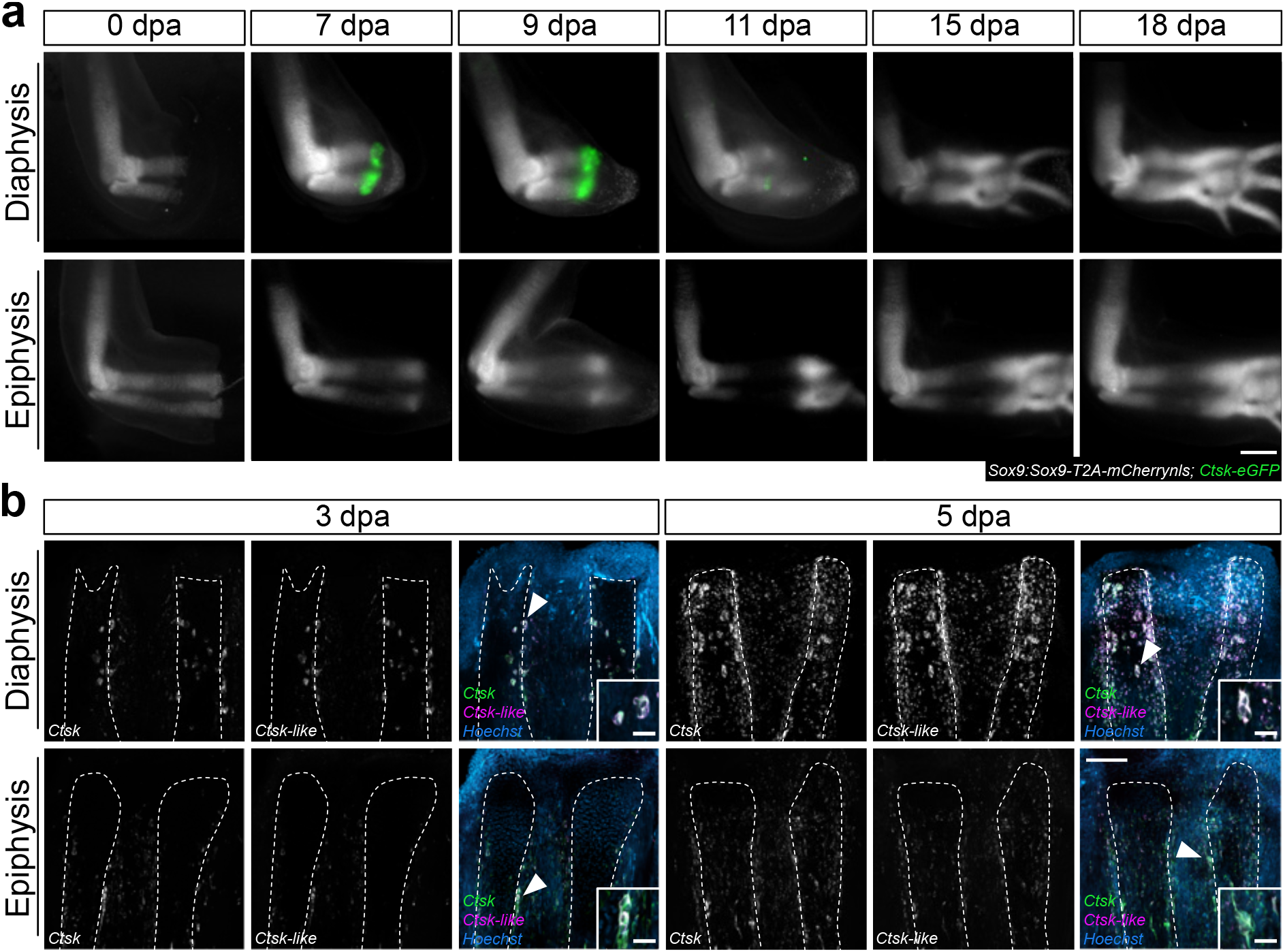
Diaphysis amputations specifically trigger osteoclast recruitment in axolotl limb regeneration. **a** Time course of osteoclast presence in diaphysis and epiphysis amputations in *Sox9::Ctsk* limbs at 0, 7, 9, 11, 15, and 18 dpa. Chondrocytes are labelled in white, osteoclasts in green. Scale bar: 1 mm. Representative images of an experiment with n = 5 animals per time point and amputation plane. **b** Hybridization chain reaction (HCR) for *Ctsk* (green) and *Ctsk-like* (magenta) at 3 and 5 dpa in representative diaphysis- and epiphysis-amputated limbs. White arrowheads indicate the multinucleated osteoclasts depicted in corresponding insets. HCR was performed in n = 3 limbs from three different animals per time point and amputation plane, and independently repeated 3 times. Scale bar: 500 µm, scale bar in insets: 50 µm.

Finally, we assessed the endogenous expression of the osteoclast marker *Ctsk* by Hybridization Chain Reaction (HCR) to determine when osteoclast presence was first differentially established in diaphysis and epiphysis amputations. While examining the *Ctsk* nucleotide sequence, we found that another gene, *Loc138578972*, was present in an adjacent region and annotated as *Ctsk-like* (Supplementary Fig. 2a). The predicted nucleotide coding sequence of this gene was 73.6% identical to the one of *Ctsk* (Supplementary Fig. 2b). Moreover, the predicted protein sequence of *Loc138578972* shared 73.4% and 73.7% sequence identity with the axolotl Ctsk and human CTSK peptide, respectively (Supplementary Fig. 2c-f). We thus concluded that the axolotl genome contains at least one additional *Ctsk*-related gene and, in keeping with the most current axolotl annotation (UKY_AmexF1_1, GCF_040938575.1), the gene annotated as *Ctsk* will continue to be referred to as *“Ctsk”*, whereas the gene *Loc138578972* will be referred to as *“Ctsk-like”* for the remainder of this work.

Analysis of both *Ctsk* and *Ctsk-like* expression showed no great difference in the number or appearance of *Ctsk*-expressing cells between diaphysis and epiphysis amputations at 1 dpa (Supplementary Fig. 3a), with the osteoclasts mainly present along the diaphyseal regions in both conditions (Supplementary Fig. 3a’ and a’’). However, by 3 dpa, large multinucleated *Ctsk*-positive cells, which are the hallmarks of mature osteoclasts, were almost exclusively associated to diaphysis amputations, and were seen in even larger numbers at 5 dpa (Fig. 2b, white arrowheads; Fig. 2b, insets). *Ctsk-like* exhibited a similar expression pattern and was largely co-expressed with *Ctsk* at all analyzed time points (Fig. 2b, Supplementary Fig. 3a, b).

Altogether, our results show that extensive osteoclast-mediated skeletal tissue resorption is a tissue-dependent event, being activated by amputations directly affecting, or in close proximity to, the calcified diaphyseal regions of the skeletal elements. Moreover, osteoclast recruitment and/or differentiation starts early within regeneration, with *Ctsk-*/*Ctsk-like*-positive large multinucleated osteoclasts appearing in diaphysis amputations as early as 3 dpa.

### Systemic and local calcium do not impact osteoclast presence

One major difference between diaphysis and epiphysis amputations is the damage to the sheath of calcified tissue surrounding the skeletal elements^7^ (Fig. 1a). In vertebrates, plasma variations in calcium trigger osteoclast recruitment into the bone so that systemic calcium concentrations are kept within homeostatic levels^13^. Moreover, it has been reported that hypocalcemia is a common occurrence in trauma patients^14,15^. This led us to hypothesize that amputations in the axolotl limb, especially diaphysis amputations, could have an effect in plasma calcium concentrations. Thus, we explored the role of systemic and local calcium in the activation of osteoclast-mediated skeletal resorption.

At the systemic level, blood plasma measurements in intact animals were consistent with previously reported values^16^. While we observed some variation in calcium concentrations between intact and amputated individuals particularly in early time points, no significant difference between conditions was observed (Fig. 3a).

**Fig. 3.**
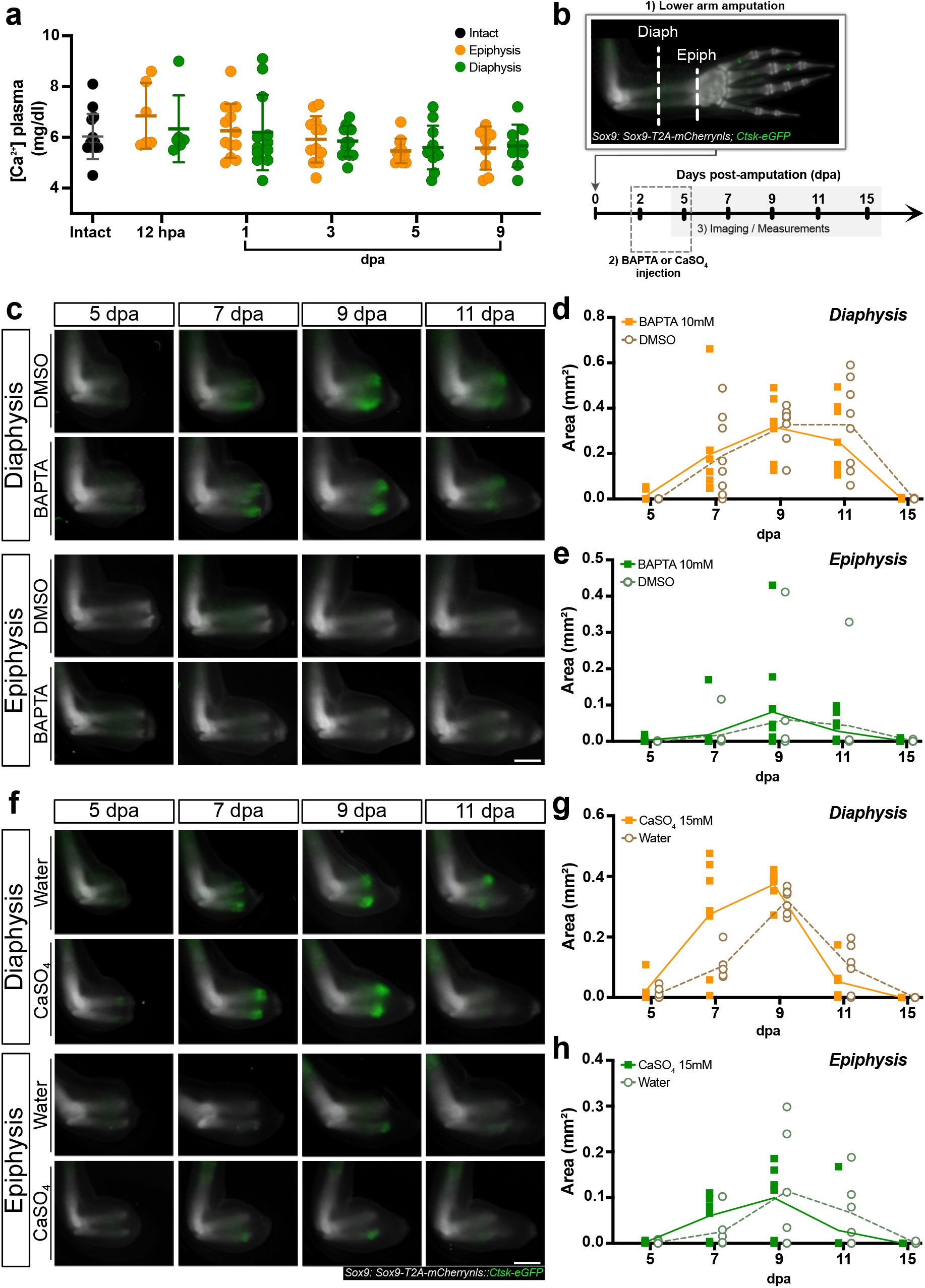
Osteoclast presence is not correlated with systemic calcium levels or impacted by local calcium changes. **a** Quantification of calcium levels in the serum of diaphysis- and epiphysis-amputated animals in intact conditions, and at 12 hpa, 1, 3, 5, and 9 dpa. The graph depicts the mean and ±sd of the combined results of 4 independent experiments using a minimum of 5 animals per time point and amputation plane. **b** Schematic representation of the experimental set up for the injections in diaphysis- and epiphysis-amputated limbs in *Sox9::Ctsk* animals. **c** Time course of osteoclast presence in representative diaphysis- and epiphysis-amputated limbs injected with DMSO (control) or 10mM BAPTA at 5, 7, 9 and 11 dpa. Scale bar: 1 mm. **d** Quantification of the area of Ctsk^+^ signal in BAPTA and DMSO injected animals in diaphysis amputations. **e** Quantification of the area of Ctsk^+^ signal in BAPTA and DMSO injected animals in epiphysis amputations. **f** Time course of osteoclast presence in representative diaphysis- and epiphysis-amputated limbs injected with water (control) or CaSO4 at 5, 7, 9 and 11 dpa. Scale bar: 1 mm. **g** Quantification of the area of Ctsk^+^ signal in water or CaSO4 injected animals in diaphysis amputations. **h** Quantification of the area of Ctsk^+^ signal in water or CaSO4 injected animals in epiphysis amputations. c and f show representative images of an experiment with n = 4 animals per time point and amputation plane. For c-h, the 2 limbs of 4 animals were injected and assessed per condition over time. For d, e, g and h, the lines show mean values over time. Source data for d, e, g, and h are provided as a Source Data file.

We next investigated if, instead, local changes in calcium levels could have a role in osteoclast recruitment and/or differentiation. We thus injected amputated limbs of *Sox9::Ctsk* animals with high concentrations of either BAPTA or CaSO4 in order to greatly decrease or increase extracellular calcium levels, respectively, locally in the tissue. We then followed osteoclast dynamics after diaphysis and epiphysis amputations (Fig. 3b). However, neither the injections of BAPTA 10mM (Fig. 3c-e), nor of CaSO4 15mM (Fig. 3f-h), had any effect on osteoclast presence in either amputation plane.

Hence, these results indicate that systemic calcium concentrations do not significantly change due to regeneration, nor due to different amputations affecting diaphyseal or epiphyseal regions. Additionally, in our experiments, changes in local calcium levels were also not sufficient to trigger differences in osteoclast recruitment and/or differentiation into the damaged skeletal tissue, even in epiphysis amputations. Overall, this suggests that neither systemic nor local changes in extracellular calcium concentrations during regeneration are sufficient to induce significant osteoclast recruitment/differentiation. However, we cannot rule out other factors working together with either systemic or local extracellular calcium levels that could trigger this process.

### Spatial transcriptomics of diaphysis & epiphysis amputations

Our data so far suggested that the skeletal tissue itself could be involved in the osteoclast recruitment and/or differentiation observed in diaphysis-amputated limbs. Thus, we asked if gene expression differences could point us to regulatory factors underlying this process. To address this, we used spatial transcriptomics in diaphysis- and epiphysis-amputated limbs at 3 and 5 dpa (Fig. 4a) – the two time points in which differential osteoclast presence first becomes evident – to investigate differences in gene expression between these two conditions.

**Fig. 4.**
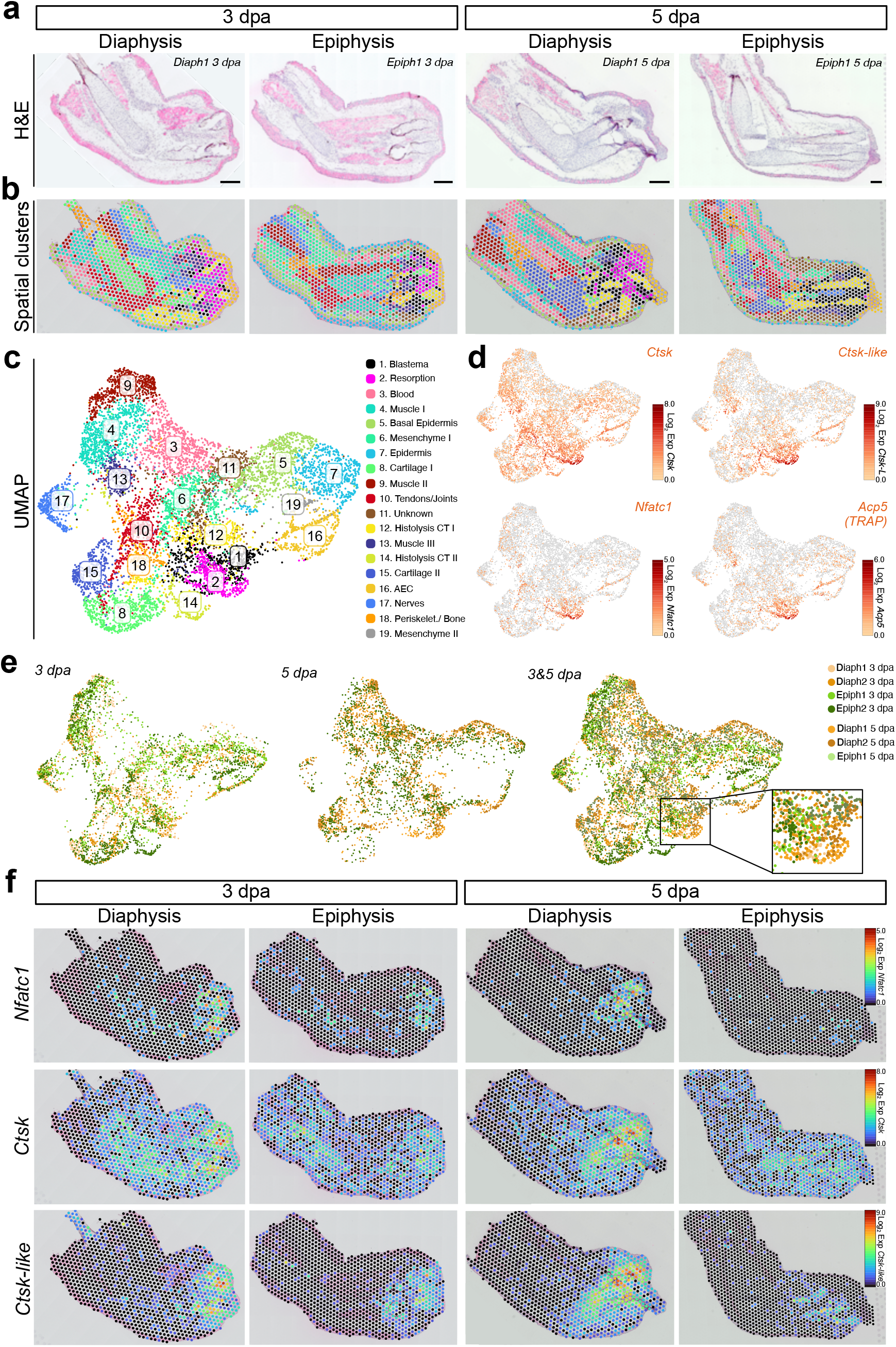
Spatial transcriptomics reveals differences in gene expression between diaphysis and epiphysis amputations. **a** Hematoxylin and Eosin (H&E) staining of representative longitudinal sections of 3 and 5 dpa diaphysis- and epiphysis-amputated limbs used for spatial transcriptomics. Representative section of n = 2 animals per time point and amputation plane, except for 5 dpa epiphysis-amputated limb, in which only one sample was used. Scale bar: 500 µm. **b** Spatial view of the 19 clusters identified by Seurat clustering analysis combining spatial dots from diaphysis- and epiphysis-amputated limbs at 3 and 5 dpa. **c** UMAP plot and cluster annotation of spatial transcriptomic dots from 3 and 5 dpa diaphysis- and epiphysis-amputated limbs. **d** UMAP plots showing the expression of the resorption-associated markers *Ctsk*, *Ctsk-like*, *Nfatc1* and *Acp5*. **e.** UMAP plot showing the contribution of each sample to the UMAP at 3 dpa (left), 5 dpa (center) and both timepoints (right). **f** Spatial expression profiles of *Nfatc1*, *Ctsk* and *Ctsk-like* in representative spatial transcriptomics sections at 3 and 5 dpa. Expression levels in d and f were calculated as Log2 fold expression. CT, connective tissue.

Clustering analysis of spatial expression dots from all samples combined revealed 19 clusters representing all major tissue types contained in the tissue sections, including epidermis (clusters 5 and 7), muscle (clusters 4, 9, and 13), cartilage (clusters 8 and 15), periskeleton/bone (cluster 18), and nerves (cluster 17) (Fig. 4b, c; Supplementary Data 2). We also detected regeneration-specific clusters, particularly a blastema cluster (cluster 1) enriched in the expression of *Kazald2*^17,18^, two clusters associated with tissue histolysis (clusters 12 and 14), and one cluster representing the AEC (cluster 16) (Fig. 4b, c; Supplementary Fig. 4a, b; Supplemental Data 2). Analysis of the expression of the resorption-associated factors *Ctsk*, *Ctsk-like*, *Nfatc1*, and *Acp5* found that these genes were highly expressed in cluster 2 (Fig. 4d; Supplementary Fig. 4b). Mapping these spots back to the tissue sections revealed that these genes were especially represented in diaphysis-amputated skeletal elements at both 3 and 5 dpa (Fig. 4e, f). Importantly, deconvolution using a previously published single-cell RNA-seq (scRNA-seq) dataset^19^ confirmed these results (Supplementary Fig. 4c; Supplementary Data 3). This, together with the fact that approximately 70% of this cluster was derived from diaphysis-amputated limbs, caused us to annotate this cluster as a resorption cluster (Fig. 4e, inset; Supplementary Fig. 4d). As expected from having a spatial dataset with supra-cellular resolution, combined with the high mobility of immune cells, we could not isolate a specific immune spatial cluster. Instead, we found immune markers spread across multiple clusters, including *Perforin1-like* (*Prf1-like*), *Proteoglycan 3* (*Prg3*), and *C-X-C motif chemokine ligand 12* (*Cxcl12*) in the mesenchyme (cluster 6), and two genes annotated as *Macrophage expressed 1-like* (*Mpeg1-like*) in both the epidermis (cluster 7) and AEC (cluster 16) (Supplementary Fig. 4b). Notably, there was an enrichment of genes related to macrophage/monocyte function like *Macrophage receptor with collagenous structure* (*Marco*), a third *Mpeg1-like* gene, *Cd68,* and *Cd14* in the resorption cluster (cluster 2) (Supplementary Fig. 4a, b). Deconvolution analysis indeed confirmed that macrophages were one of the major cell types comprising the resorption cluster (Supplementary Fig. 4b, c; Supplementary Data 3). This points at the possibility that, similarly to mammals^20,21^, these myeloid cells could act as possible sources of osteoclast progenitors during regeneration.

Taken together, we show that spatial transcriptomics is able to identify clear differences in gene expression between diaphysis and epiphysis amputations, and that a specific resorption cluster enriched in myeloid markers is predominately present in diaphysis-amputated limbs.

### RANK/RANKL likely orchestrate osteoclast-mediated resorption

In examined vertebrates, osteoclast progenitors differentiate into mature osteoclasts under the influence of *RANKL* (*Tnfsf11*) and *RANK* (*Tnfrsf11a*)^22–24^. In our spatial dataset, we found that the majority of spatial dots in the resorption cluster (cluster 2) did indeed express both genes (Fig. 5a; Supplementary Fig. 5a), with *RANK* slightly upregulated in diaphysis amputations at both time points (Fig. 5b, bottom row). In contrast, differences in *RANKL* expression were mostly observed at 5 dpa, being expressed in more spatial dots and with overall higher expression levels in diaphysis amputations (Fig. 5b, top row).

**Fig. 5.**
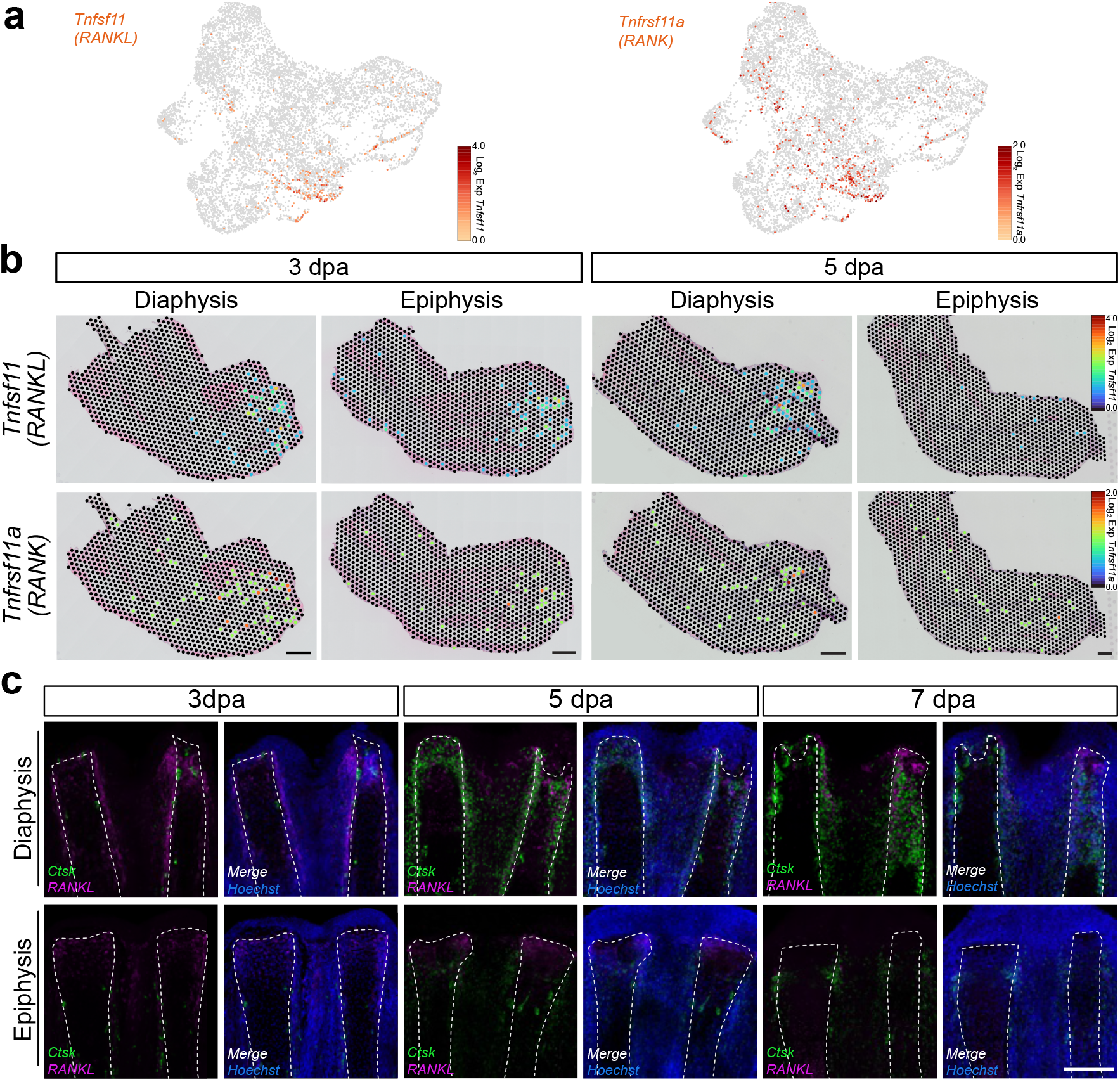
The RANK/RANKL system likely orchestrates tissue-dependent osteoclast-mediated skeletal resorption. **a** Expression of *Tnfsf11 (RANKL)* and *Tnfrsf11a (RANK)* in the UMAP from diaphysis- and epiphysis-amputated limbs at 3 and 5 dpa. Expression levels were calculated as Log2 fold expression. **b** Spatial expression profiles of *RANKL* and *RANK* in representative spatial transcriptomics sections at 3 and 5 dpa. Scale bar: 500 µm. **c** HCR for *Ctsk* (green) and *RANKL* (magenta) at 3, 5 and 7 dpa in representative diaphysis- and epiphysis-amputated limbs. HCR was performed in n = 3 limbs from three different animals per time point and amputation plane, and independently repeated 3 times. Scale bar: 300 µm.

To validate our spatial data and identify the cells expressing *RANK* and *RANKL*, we performed HCR for these two genes in diaphysis and epiphysis amputations at 3, 5, and 7 dpa (Fig. 5c). In epiphysis-amputated limbs, *RANKL* was observed primarily in the cartilage cells closest to the AEC at 3 dpa. Its expression was decreased at 5 dpa and, by 7 dpa, only very low levels of *RANKL* were detected in these limbs (Fig. 5c, bottom row). This contrasted with diaphysis-amputated limbs, in which *RANKL* was robustly expressed, especially in periskeletal cells and hypertrophic chondrocytes at all analyzed time points (Fig. 5c, top row). Furthermore, HCR staining for *RANKL* and *Ctsk* showed that *Ctsk^+^* cells were closely associated to cells expressing *RANKL* in diaphysis amputations (Fig. 5c, top row).

On the other hand, *RANK* expression was more prevalent in diaphysis-amputated limbs at 3 and 5 dpa and was frequently co-expressed with *Ctsk* (Supplementary Fig. 5b), suggesting that these cells were differentiating osteoclasts. We further validated this by assessing the expression of *Nfatc1*, the master transcriptional factor of osteoclastogenesis^25^ that promotes the expression of *Ctsk* and components of the vacuolar V-ATPase^26^. Indeed, we found that *RANK*-expressing cells co-expressed *Nfatc1*, especially in diaphysis-amputated limbs (Supplementary Fig. 5c). Interestingly, in these limbs, we also observed cells that co-expressed *RANK* and *Nfatc1*, but not *Ctsk*, suggesting that these could be osteoclast progenitors undergoing differentiation (Supplementary Fig. 5c, white arrowheads). Finally, *RANK^+^* cells were found in close proximity to *RANKL^+^* areas (Fig. 5c; Supplementary Fig. 5b, c). This explained why these two factors, which represent a ligand-receptor pair typically present in different cell types^9,23^, appear in the same spatial cluster.

Overall, our data indicates that *RANKL* expression is upregulated and sustained in periskeletal cells and hypertrophic chondrocytes upon calcified diaphysis amputations, which may drive the differentiation of *RANK^+^*/*Nfatc1*^+^ precursors into mature osteoclasts. In contrast, in epiphysis amputations, *RANKL* is only transiently expressed in distal chondrocytes. This suggests that, similar to mammalian osteoclastogenesis, the RANK/RANKL system has a key role in osteoclast differentiation during axolotl regeneration, and that the differential activation of *RANKL* expression after injury to the calcified diaphyseal region likely triggers tissue-dependent skeletal resorption.

#### Loc138491483/Ccl24-like induces ectopic osteoclast presence

As osteoclast-mediated tissue resorption occurs in a relatively short and well-defined time window during limb regeneration, we next searched for regeneration-specific factors that could regulate this process.

For this, we reasoned that osteoclast progenitors and/or immature osteoclasts, identified by *Nfatc1* expression, would be in close spatial proximity to any putative signal factor at 3 and 5 dpa. Our analysis revealed that the top 15 differentially expressed genes in *Nfatc1*-enriched spatial spots (Log2 Expression > 2) were enriched in genes heavily involved in osteoclast function, such as V-ATPase subunit genes (*Atp6v0c*, *Atp6v0d2*, *Atp6v1b2*) and MMPs/ECM remodeling genes (*Ctsk*, *Ctsk-like*, *Mmp9*) (Fig. 6a; Supplementary Data 4). However, there was one gene, *Loc138491483* (hereafter referred to as *Loc483*), that was seemingly not directly related to osteoclast function while still being associated with the resorption cluster (cluster 2) (Fig. 6b, c) and to the skeletal tissue in diaphysis amputations (Fig. 6d). We then validated these findings by HCR at an earlier time point (1 dpa) and at 3 and 5 dpa. While no differences were observed in the expression of this gene at 1 dpa, *Loc483* became highly expressed in diaphysis-amputated limbs at both 3 and 5 dpa when compared to epiphysis-amputated limbs (Fig. 6e). We then asked if *Loc483* was expressed in the osteoclast progenitors themselves, or by the surrounding niche to recruit them. HCRs for *Loc483* and *Ctsk* at 1, 3 and 5 dpa showed that both genes only very partially overlap, which would more support the latter hypothesis (Supplementary Fig. 6a). Yet, when we assessed the co-expression of *Loc483* and *Nfatc1*, we found that the great majority of *Loc483*^+^ cells were also *Nfatc1^+^*. This suggests that *Loc483* is predominantly present in osteoclast progenitors, with only some maturing osteoclasts continuing to express it (Supplementary Fig. 6b, b’’, white arrowhead). However, *Nfatc1* was not restricted to *Loc483*^+^ cells (Supplementary Fig. 6b-b’’, outlined arrowhead), indicating that either not all osteoclast progenitors are *Loc483^+^* or, alternatively, that these *Loc483^-^/Nfatc1^+^* cells are not part of the osteoclast lineage, as previously reported in mammals^27^.

**Fig. 6.**
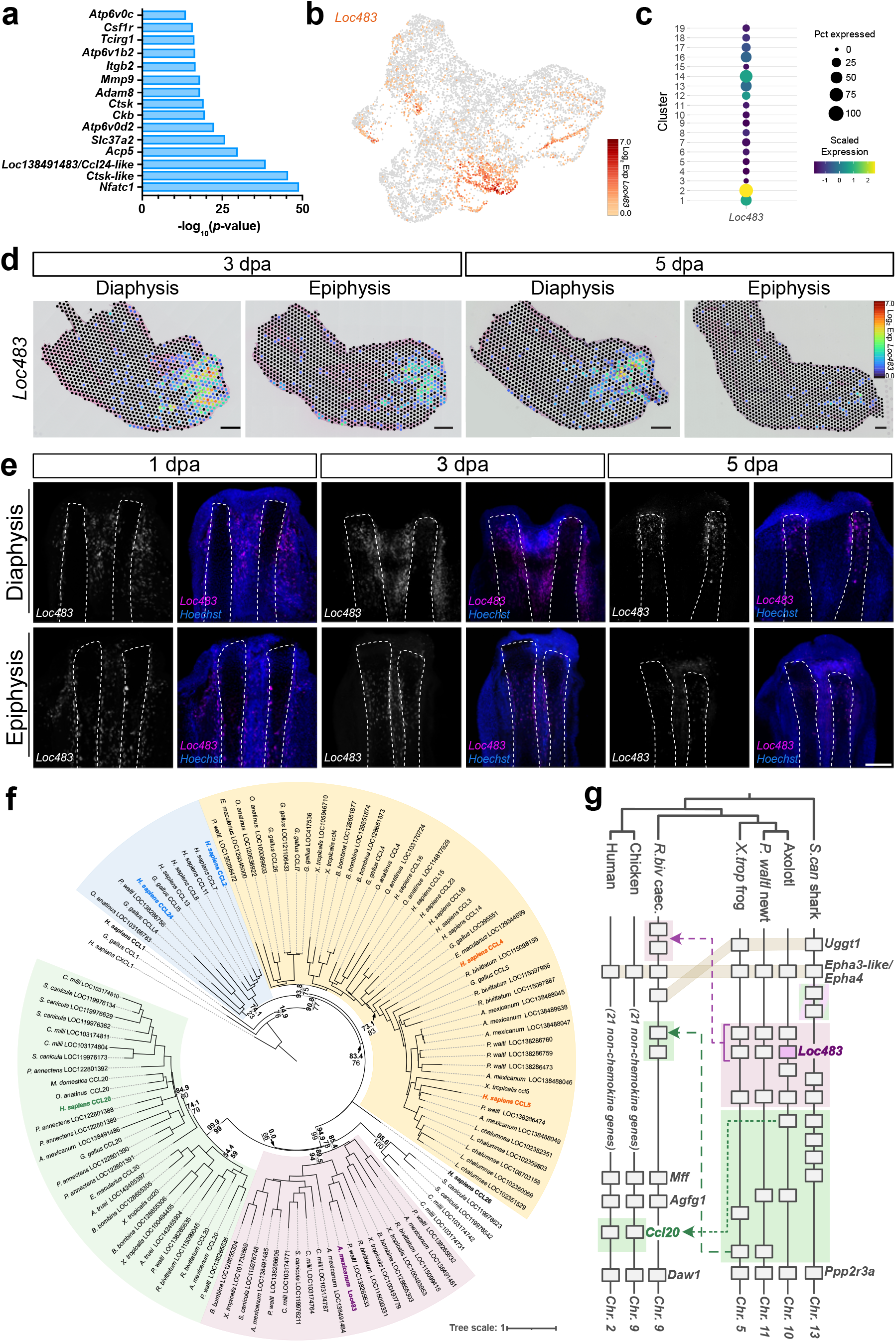
*Loc483* is upregulated in diaphysis amputations and is part of a clade of chemokines maintained only in amphibians and sharks. **a** Top 15 differentially expressed genes in *Nfatc1*-expressing spatial transcriptomics spots (exact negative binomial test, *p*-values adjusted for multiple testing using the Benjamini-Hochberg procedure). **b** Expression of *Loc483* in the UMAP from diaphysis- and epiphysis-amputated limbs at 3 and 5 dpa. **c** Dot plot showing expression of *Loc483* in the 19 annotated spatial transcriptomics clusters of diaphysis- and epiphysis-amputated limbs at 3 and 5 dpa. **d** Spatial expression profiles of *Loc483* in representative spatial transcriptomics sections at 3 and 5 dpa. Scale bar: 500 µm. **e** HCR for *Loc483* at 1, 3 and 5 dpa in diaphysis- and epiphysis-amputated limbs. HCR was performed in n = 3 limbs from three different animals per time point and independently repeated 4 times. Scale bar: 300 µm. **f** Maximum-likelihood (ML) phylogenetic tree of chemokines selected based on syntenic similarities to axolotl *Loc483* and human *CCL2*, *CCL4*, and *CCL24*. Support values displayed for key nodes, with SH-aLRT^80^ values on top in bold and ultrafast bootstrap^81^ approximation values on the bottom. Clades colored based on grouping with specific chemokines: Blue – human *CCL2* & *CCL24*, Yellow – human *CCL4* & *CCL5*, Pink – axolotl *Loc483*, Green – human *CCL20*. Scale bar corresponds to the mean number of amino acid substitutions per site. **g** Schematic representation of genomic regions containing Loc483- and Ccl20-related chemokines across several vertebrates. Non-chemokine genes *Uggt1* and *Epha3-like*/*Epha4* (name varying between species) were used as anchor genes of the region. Colors indicate clades of the ML tree. Green-pink gradient shading of two shark chemokines represents shark specific outgroup to Loc483 & Ccl20 clades (not highlighted in ML tree). Orthologous chemokines are aligned when possible, and otherwise marked with dashed arrows. Gene size and distances between genes not drawn to scale. *S. can*, *Scyliorhinus canicula*, *P. waltl*, *Pleurodeles waltl*; *X. trop*, *Xenopus tropicalis*; *R. biv*, *Rhinatrema bivittatum*; caec., caecilian.

To clarify this point, we investigated *Loc483*^+^ cells in the scRNA-seq dataset of regenerating limbs previously used for deconvolution^19^. This confirmed that *Nfatc1* and *Loc483* expression considerably overlapped (Supplementary Fig. 6c), supporting the HCR findings. However, this also uncovered the existence of *Loc483*^+^ cells that did not express *Nfatc1* (Supplementary Fig. 6d, e). Furthermore, co-expression analysis of *Loc483* and *RANK* indicated that only a few cells expressed both genes (Supplementary Fig. 6f). This all suggests that only a subset of osteoclast-lineage cells express *Loc483*, the majority of them most likely being progenitors. However, other cell types throughout the niche appear to also express this gene.

We next sought to determine the nature of the *Loc483* gene. While it is annotated in the axolotl genome as *C-C motif chemokine 24-like* (*Ccl24-like*), an initial phylogenetic search via the webserver aLeaves^28^ revealed that its most closely related protein sequences were *Monocyte Chemoattractant Protein-1/C-C motif chemokine 2* (*Mcp1/Ccl2*) in the caecilian *Rhinatrema bivittatum* and *C-C motif chemokine 4* (*ccl4*) in the frog *Xenopus tropicalis*. Additional matches could only be found in two species of sharks and in the paddlefish, suggesting that it was not orthologous to human *CCL24*, but providing little clue to its true relationship with other chemokines. Thus, we performed an extensive molecular phylogenetic analysis using chemokines from a diverse collection of species, revealing that *Loc483* is part of a distinct clade of chemokines maintained only within amphibians and cartilaginous fish. Both maximum likelihood and Bayesian inference trees revealed this clade to be most related to that of *Ccl20* (Fig. 6f; Supplementary Fig. 7b), with the two clades together forming a clear outgroup to the clades containing *Ccl24*, *Ccl2*, and *Ccl4*. This was strongly supported by synteny analysis, which revealed members of the *Loc483* clade to be colinear with members of the *Ccl20* clade in sharks and most amphibians (Fig. 6g), suggesting an origin via tandem duplication. Meanwhile, *Ccl24*, *Ccl2*, and *Ccl4* were located on entirely different chromosomes (Supplementary Fig. 7a). Thus, our work demonstrates that *Loc483* is part of an ancient group of chemokines which originated in the last common jawed vertebrate ancestor but has been lost in all tetrapods except amphibians.

As a chemokine, *Loc483* was a potential chemoattractant for macrophages and monocytes, both previously reported as sources of osteoclast progenitors^20,21,29^. Thus, to test if *Loc483* could promote osteoclast recruitment and/or differentiation, we overexpressed this gene in *Sox9::Ctsk* animals. For that, we co-electroporated limbs with a construct containing the coding sequence of *Loc483* under the control of the ubiquitous promoter CAGGS (*CAGGS-Loc483*), together with a reporter plasmid containing mCherry driven by the same promoter (*CAGGS-mCherry)* (Fig. 7a, Supplementary Fig. 8). The contralateral blastema was electroporated only with the *CAGGS-mCherry* construct as a control (Fig. 7a, Supplementary Fig. 8). We found that, in diaphysis amputations, overexpression of *Loc483* had no effect on the presence of osteoclasts (Fig. 7b, c). However, *Loc483* was able to ectopically induce the presence of osteoclasts in epiphysis-amputated limbs (Fig. 7b, d).

**Fig. 7.**
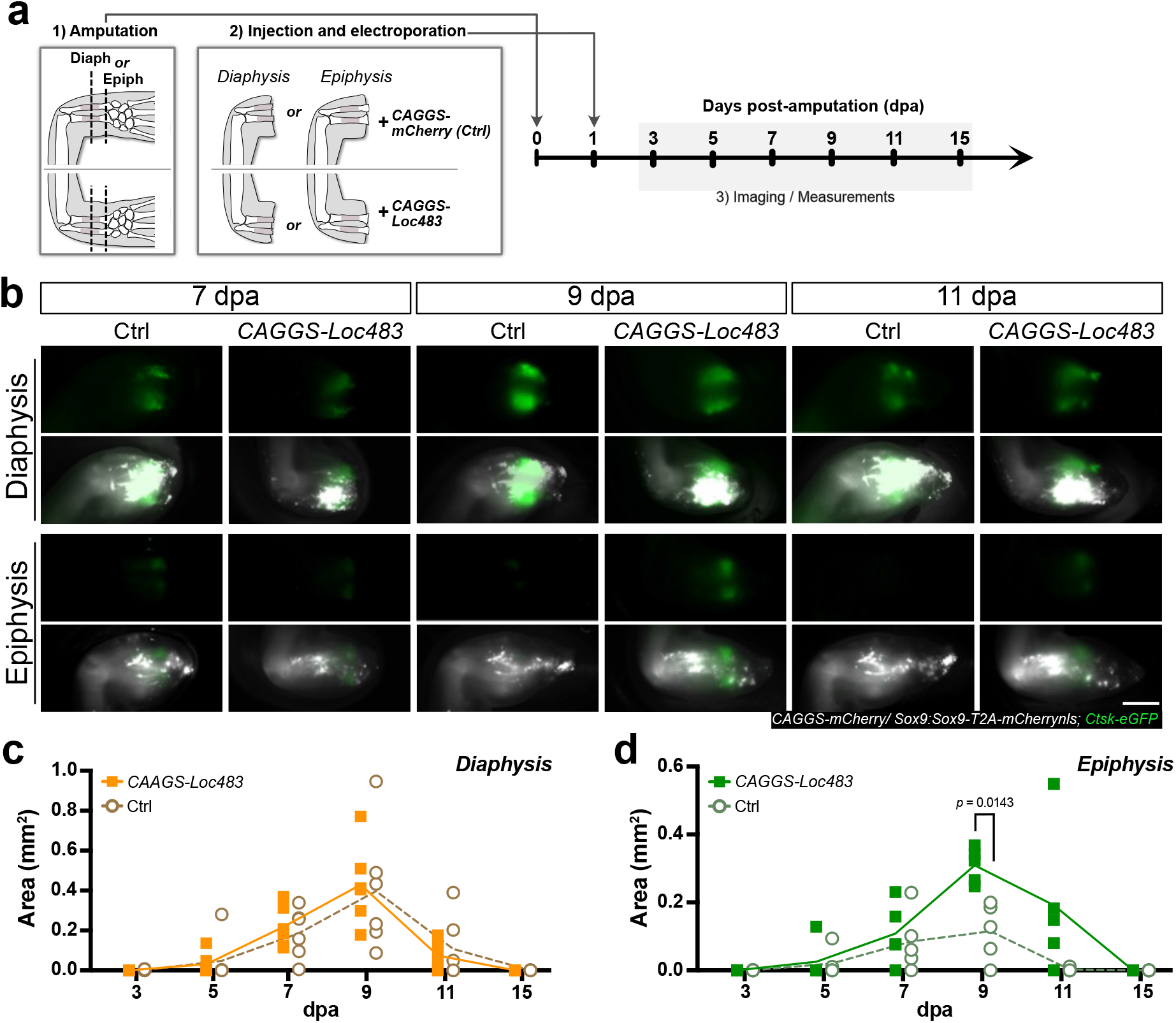
*Loc483* is sufficient to ectopically recruit osteoclasts into epiphysis-amputated limbs. **a** Schematic representation of the experimental set up used for the electroporation of *Loc483* in diaphysis- and epiphysis-amputated limbs of *Sox9::Ctsk* animals. Images of limbs were created in Affinity Designer 2. **b** Time course of osteoclast presence in representative diaphysis- and epiphysis-amputated limbs injected and electroporated with control plasmid (*CAGGS-mCherry*) or control plasmid + *CAGGS-Loc483* at 7, 9, and 11 dpa. Representative images of an experiment with n = 3 animals per time point and amputation plane. Scale bar: 1 mm. **c** Quantification of the area of Ctsk^+^ signal in *CAGGS-Loc483* and control (Ctrl) animals in diaphysis amputations. **d** Quantification of the area of Ctsk^+^ signal in *CAGGS-Loc483* and control (Ctrl) animals in epiphysis amputations. For c and d, the graphs represent the combined results of 2 independent experiments using a total of 5 limbs of 5 different animals per condition. The lines show mean values over time ± sd. Statistical significance for *CAGGS-Loc483* vs Ctrl electroporated animals was assessed by a two-way ANOVA with Tukey’s post-hoc test. Source data for c and d are provided as a Source Data file.

Thus, these results show that *Loc483* is a chemokine within the amphibian immune system, and that its expression is sufficient to recruit and/or differentiate osteoclasts to the amputation site.

### Diaphysis amputations recruit osteoclast to epiphyses

We then investigated if recruitment of osteoclasts is specific to the injured calcified tissue. For that, we performed staggered amputations in the radius and the ulna of *Sox9::Ctsk* animals. In these limbs, the ulna of the left limb was amputated at the level of the epiphysis, while its radius was amputated through the diaphysis. The right limb of the same animal was amputated in the reverse way, and animals were analyzed at 9 dpa (Fig. 8a). We found that, in the majority of the limbs, an amputation in the diaphysis was sufficient to recruit *Ctsk^+^* cells into the neighboring epiphysis-amputated skeletal element (Fig. 8b-d). However, in approximately 35% of the cases, resorption remained restricted to the diaphysis-amputated limb (Fig. 8b, An#14). Interestingly, we found that both scenarios could occur in the same animal (Fig. 8b, An#13).

**Fig. 8.**
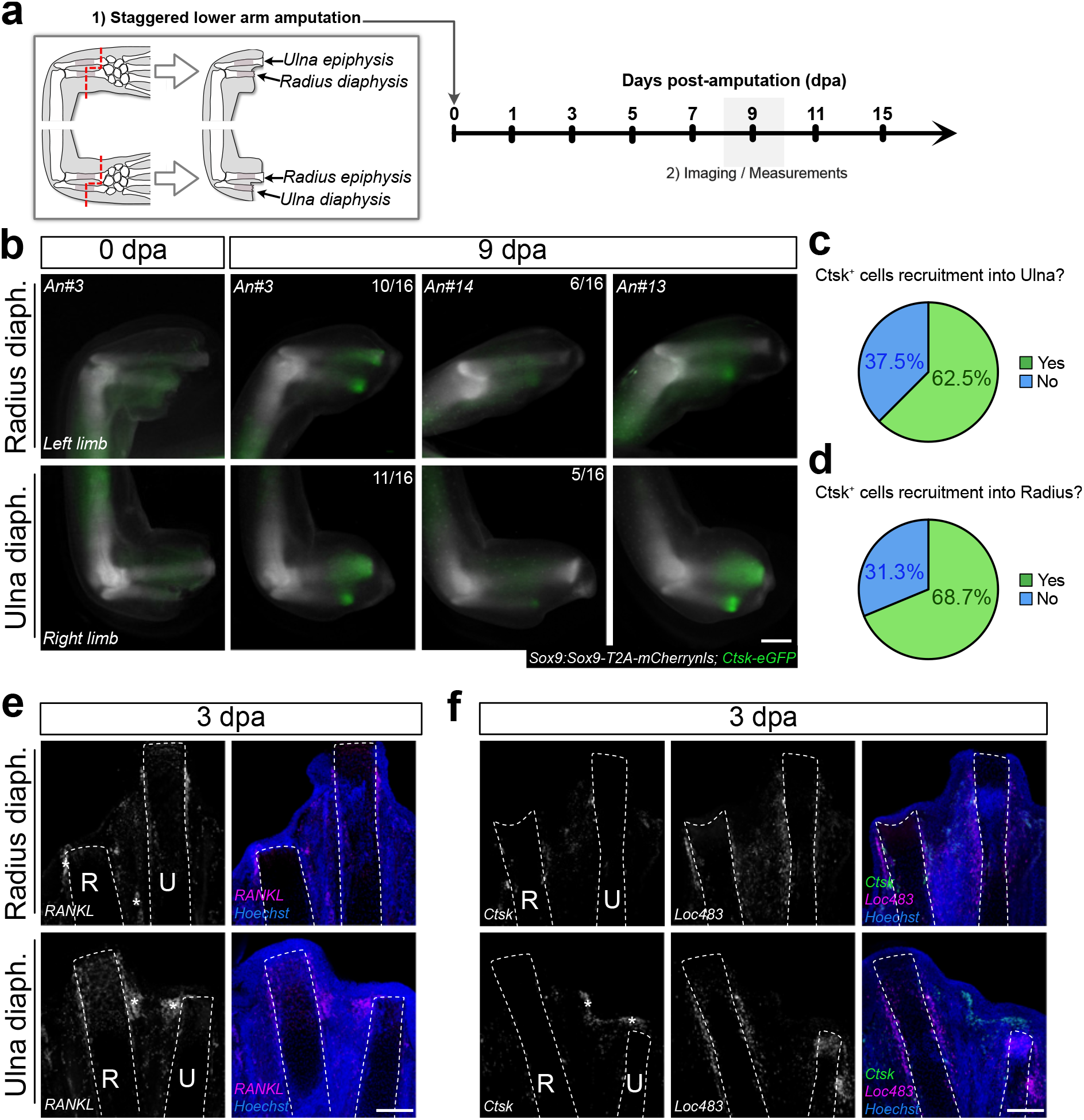
Diaphysis amputations can induce osteoclast recruitment into epiphysis-amputated skeletal elements. **a** Schematic representation of the experimental set up used for staggered amputations in *Sox9::Ctsk* animals. In each animal, the radii of left limbs were amputated in the diaphysis, whereas ulnas were amputated in epiphyseal regions. The converse amputations were performed in the right limb. Images of limbs were created in Affinity Designer 2.**b** Representative images of left and right limbs of three *Sox9::Ctsk* animals at 0 dpa and 9 dpa. Animal#3 (An#3) is showed at 0 dpa and at 9 dpa to illustrate limb morphology and Ctsk^+^ signal in the two time points. The number of limbs with the same pattern of Ctsk^+^ signal out of the total number of limbs (n = 16) is indicated in the top right corner. Scale bar: 1 mm. **c** Percentage of epiphysis-amputated ulnas in limbs with diaphysis-amputated radii exhibiting Ctsk^+^ cell recruitment. **d** Percentage of epiphysis-amputated radii in limbs with diaphysis-amputated ulnas exhibiting Ctsk^+^ cell recruitment. Data in b-d represents the combined results of 2 independent experiments using a total of 16 animals (16 left limbs and 16 right limbs). **e** HCR for *RANKL* in limbs with diaphysis-amputated radii or diaphysis-amputated ulnas at 3 dpa. HCR was performed once in n = 3 limbs from different animals per time point and amputation plane. Scale bar: 300 µm. **f** HCR for *Ctsk* (green) and *Loc483* (magenta) in limbs with diaphysis-amputated radii or diaphysis-amputated ulnas at 3 dpa. HCR was performed once in n = 3 limbs from different animals per time point and amputation plane. R, radius, U, ulna. Scale bar: 300 µm.

Expression analyses for *RANKL* and *Loc483* at 3 dpa confirmed that these genes were present in diaphysis-amputated skeletal elements (Fig. 8e, f), as previously observed in non-staggered amputations. However, they were also strongly expressed in the skeletal element amputated in the epiphysis, which correlated with the presence of osteoclasts in the two skeletal elements.

In conclusion, our data suggests that amputations in the calcified diaphysis trigger the molecular pathways associated with osteoclast recruitment, which then mobilize these cells to the skeletal elements regardless of the amputation plane.

### Tissue composition at the amputation plane affects the AEC

Our spatial transcriptomic data accurately captured local differential gene expression in diaphysis amputations when compared to epiphysis amputations. However, this technique is limited to the areas captured on the slide. To complement the spatial information of our datasets and get an overview of gene expression in diaphysis- and epiphysis-amputated limbs, we performed bulk RNA sequencing (RNA-seq) in intact diaphysis and epiphysis regions, and in the distal-most region of regenerating limbs (Supplementary Data 5; Supplementary Fig. 9). This found 100 and 80 differentially expressed genes (DEGs) between the two amputation planes at 3 and 5 dpa, respectively (Supplementary Data 6, 7).

Unexpectedly, genes related to bone resorption and osteoclast function such as *Ctsk*, *Ctsk-like*, *Nfatc1*, and *Acp5*, although upregulated after either type of amputation compared to intact conditions, were not differentially expressed between diaphysis- and epiphysis-amputated limbs at either 3 or 5 dpa in our bulk RNA-seq dataset (Supplementary Fig. 10a-e). This was especially surprising for the first three, given that our spatial transcriptomics and subsequent validation by HCR staining showed a clear higher expression of these genes in diaphysis amputations as early as 3 dpa (Fig. 2b; Fig. 4f; Supplementary Fig. 4b).

One possible explanation for this discrepancy is that cells from areas not captured with these techniques are expressing these genes, masking the local change observed in individual tissue sections. Alternatively, if these genes are being utilized for processes unrelated to osteoclastogenesis^22,27,30–32^ and their expression varies independently to it, there may exist substantial variability in expression that is not impacted by the amputation plane. Variability could also be artificially caused by the susceptibility of bulk RNA-seq data to differences in the composition of collected sample, which can skew the expression levels of genes that are limited to specific regions of the tissue. Such variability can be observed in the case of one of the 3 dpa epiphysis-amputated limbs (Supplementary Fig. 9), although we cannot confirm what the underlying reason may be. Regardless, the high variability would reduce the statistical power, potentially causing these genes to not be differentially expressed. Such a possibility highlights the benefits of our multiple methods of validation.

Many of the DEGs detected by bulk RNA-seq, were either uncharacterized genes (i.e., protein-coding genes with no further annotation) or predicted to be non-coding RNAs (ncRNAs). Indeed, a look into the top 15 to 20 upregulated DEGs in diaphysis and epiphysis amputations at the two time points showed that 30% to 60% of these genes were computationally annotated as ncRNAs (Supplementary Fig.10a-d). Consequently, we filtered out both ncRNAs and uncharacterized genes for all further downstream analyses (Supplementary Data 8). This resulted in 60 and 35 upregulated protein-coding DEGs in epiphysis amputations at 3 and 5 dpa, respectively (Supplementary Fig. 10a-b), while diaphysis amputations exhibited only 10 and 8 upregulated protein-coding DEGs at these time points, respectively (Supplementary Fig. 10c-d).

Gene ontology (GO) analysis for biological processes with these genes (Supplementary Data 9) showed that upregulated DEGs in 3 dpa epiphysis amputations were enriched in terms associated with response against viruses and with protein folding (Supplementary Fig. 10f). In contrast, upregulated DEGs in 3 dpa diaphysis amputations were mostly associated with muscle tissue, as we saw *Myl4*, *Loc138582818/Myh4-like*, and *Loc138573356/Tpm1-like* representing 3 out of the 10 upregulated DEGs (Supplementary Fig.10 c, f). However, this could likely represent differences in the amount of muscle present in epiphyseal and diaphyseal regions^33^. Importantly, *Loc483* was also significantly upregulated in diaphysis amputations at 3 dpa (Supplementary Fig. 10c, g), which agreed with our spatial transcriptomics datasets, as well as our previous expression and functional validations. As for at 5 dpa, DEGs identified in epiphysis-amputated limbs were significantly enriched for GO terms associated with epidermis and with ECM production, whereas only *Cell adhesion* (GO:0007155) was found to be significantly associated to diaphysis-amputated limbs at the same time point (Supplementary Fig. 10f).

When we performed GO analysis for cellular component (CC) and molecular function (MF) instead (Supplementary Data 9), we saw that the DEGs found in diaphysis-amputated limbs at 3 and 5 dpa were also significantly enriched in ECM and muscle-related CC terms (Fig. 9a). Moreover, MF terms enriched for these amputations at 3 dpa were generally in agreement with the CC analysis, as they were similarly associated to muscle function, although no MF term was significantly enriched at 5 dpa. In contrast, DEGs in 3 and 5 dpa epiphysis-amputated limbs were enriched for CC terms associated to secretion or extracellular space, as well as to epidermis. Meanwhile, enriched MF terms in this amputation plane at 3 dpa were associated to protein folding, acetylcholine receptor signaling, and exosome function (Fig. 9a). Interestingly, by 5 dpa, these MF terms changed to being enriched in protease inhibitor-related ones, such as *Serine protease inhibitor* (KW-0722) and *Serine-type endopeptidase inhibitor activity* (GO:0004867).

**Fig. 9.**
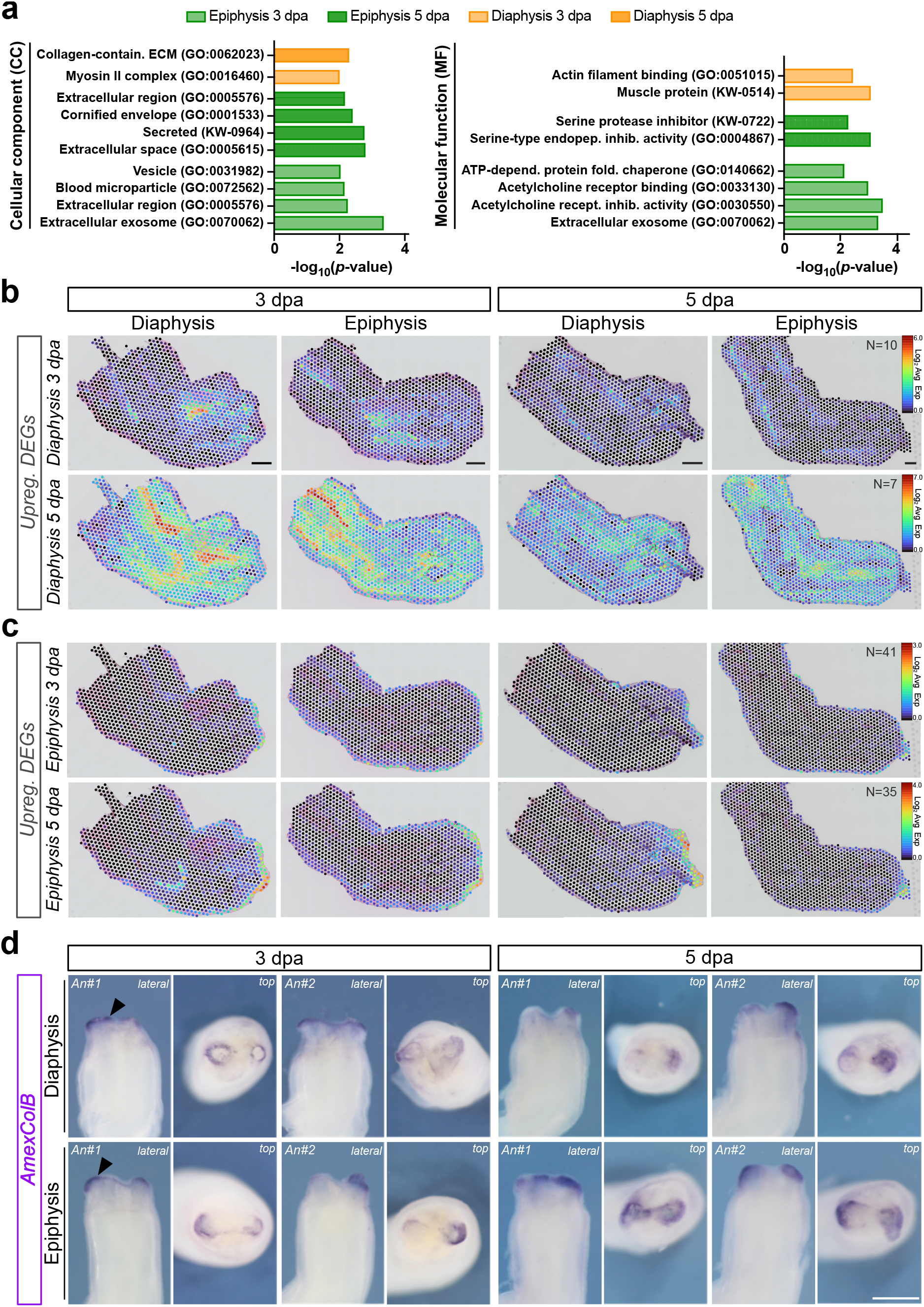
Diaphysis and epiphysis amputations induce changes in the transcriptomic profile of their AEC. **a** GO analysis for Cellular Component (CC) and Molecular Function (MF) terms in upregulated DEGs found in bulk RNA-seq of diaphysis- and epiphysis-amputated limbs at 3 and 5 dpa (*p* < 0.01, Fisher’s Exact test). **b** Spatial transcriptomic profile of the average expression of upregulated bulk RNA-seq DEGs in diaphysis-amputated limbs at 3 dpa (10 genes) and 5 dpa (7 genes). Scale bar: 500 µm. **c** Spatial transcriptomic profile of the average expression of upregulated bulk RNA-seq DEGs in epiphysis-amputated limbs at 3 dpa (41 genes) and 5 dpa (35 genes). In b and c, expression levels were calculated as Log2 average expression. **d** Whole-mount in situ hybridization (WISH) of *AmexColB* in diaphysis- and epiphysis-amputated limbs at 3 and 5 dpa. Two animals per time point and condition are displayed in lateral and top view. Scale bar: 1 mm. WISH was performed in n = 3 limbs from three different animals per time point and amputation plane, and independently repeated 3 times.

Given the prevalence of GO terms related to epidermis and secretion enriched in upregulated DEGs in the bulk RNA-seq from epiphysis-amputated limbs, we hypothesized that these genes were mostly being expressed in the AEC. To explore this possibility, we leveraged our spatial transcriptomics dataset to localize the expression of these DEGs within the context of this tissue. For that, we first took all DEGs from the bulk RNA-seq previously used for GO analysis, removed genes that were not expressed or were very lowly expressed in the spatial transcriptomics dataset (Log2 Expression <1). We then used Loupe Browser to display the combined average expression levels of upregulated DEGs at 3 and 5 dpa of epiphysis (N= 41 and 35 genes, respectively) and diaphysis amputations (N= 10 and 7 genes, respectively) in our spatial tissue sections (Fig. 9b, c). This approach revealed that, on average, upregulated DEGs found by bulk RNA-seq in epiphysis amputations at 3 and 5 dpa (Fig. 9c) tended to be expressed in the AEC. Indeed, visualization of representative upregulated DEGs in epiphysis-amputated limbs at 3 (*Psat1*) and 5 dpa (*Klf17* and *Csdn*), found that the spatial dots with the highest expression levels were localized to the AEC, with variable numbers of low-expressing dots scattered throughout the inner limb tissue (Supplementary Fig.11a). This was also observed when the average expression of DEGs annotated as ncRNAs was analyzed (Supplementary Fig.11c). However, no such trend was detected with upregulated DEGs in diaphysis limbs at either time point (Fig. 9b, Supplementary Fig.11b). The combined average expression of upregulated DEGs in this amputation plane was instead found throughout the limb tissue, which included the epidermis and AEC (*Msln*), connective tissue (*Lum*, *Postn*), muscle (*Myl4*, *Myh4-like, Tpm1-like*), nerves (*Mpz*), and tendons/joints (*Thbs4*) (Supplementary Fig. 12a).

Surprisingly, the combined average expression of genes in the spatial transcriptomics sections did not seem to reflect the differences in expression of DEGs found by bulk RNA-seq. In particular, DEGs upregulated in epiphysis amputations at 5 dpa identified by bulk RNA-seq did not appear to be altered in spatial transcriptomics compared to diaphysis amputations. This could be explained as either resulting from the flattening of differences between diaphysis and epiphysis amputations caused by averaging the expression of multiple genes with different expression levels, by saturation of the spatial dots with highly expressed genes, or simply by the fact that the analyzed tissue sections did not contain the particular region of the AEC where gene expression of these DEGs was at its strongest.

We then decided to further investigate these transcriptomic changes in the AEC by visualizing the expression of *AmexColB*, an amphibian-specific colagenase^34^ found to be differentially expressed in the bulk RNA-seq datasets of epiphysis-amputated limbs (Supplementary Fig. 10b; Supplementary Fig. 12b). Whole-mount in situ hybridization (WISH) for this gene at 3 and 5 dpa revealed that *AmexColB* was expressed in the AEC in both diaphysis- and epiphysis-amputated limbs (Fig. 9d, black arrowheads). However, the staining appeared both stronger and more widespread in epiphysis-amputated limbs, especially in a top view, which agreed with the bulk RNA-seq results (Fig. 9d).

Altogether, the transcriptomic profile differences of the AEC in diaphysis vs. epiphysis amputations suggest adaptations of its gene expression during limb regeneration that depend on the damaged tissue, particularly in its secretory profile. However, more work is still needed to clarify how the transcriptional differences observed between the AECs result in significant functional consequences, as well as what is the role of the many differentially expressed ncRNAs in limb regeneration.

## Discussion

A crucial aspect of successful limb regeneration is the robust integration between the tissues of the newly regenerated body part and the previously existing structure. Skeletal tissue integration, in particular, is especially complicated by its spatially and temporally dynamic tissue composition, in which a fully cartilaginous skeleton at the end of development becomes progressively mineralized until achieving adult patterns of ossification around sexual maturation^7^. Recently, our lab showed that the tightly-controlled clearing of skeletal tissue by osteoclasts is essential for skeletal integration^6^. However, it was unknown if osteoclast-mediated resorption promoted integration in amputation planes exposing different cell and extracellular matrix compositions. Moreover, it was also unclear how this process was regulated, especially given its fast-acting nature during regeneration that contrasts with the slow bone remodeling in fractures^35^.

Here we show that skeletal integration strategies during axolotl limb regeneration are customized to the tissues damaged by the amputation, and that these adaptations are initiated early on. We establish that osteoclast-mediated skeletal resorption is primarily activated in amputations through calcified diaphyseal regions, but not through the cartilaginous epiphysis. We also demonstrate that the expression of the chemokine *Loc483* is specifically sustained in diaphysis-amputated limbs and is sufficient to promote osteoclast presence in amputated limbs. Finally, we reveal that the transcriptomic profile of the AEC is modified by the amputation plane, suggesting that this structure may be adapted to the types of tissues injured by the amputation.

Previous studies found that successful tissue integration in the axolotl is impacted by factors like defect size^35–37^, vitamin D^38^ and positional identity incompatibility^39^. In this work, we set to investigate the importance of the skeletal composition itself at the amputation plane. To address this, we performed amputations in two sites that were closely localized within the same limb segment, and thus mainly differed on the affected skeletal tissue composition. We found that, unlike with injuries affecting cartilaginous epiphyses, amputations in the calcified diaphysis undergo extensive osteoclast recruitment and/or differentiation and skeletal tissue resorption. However, exactly how this process contributes to skeletal integration in diaphyseal amputations is still unknown. We propose two non-mutually exclusive hypothesis, which may even act synergistically.

The first hypothesis is that this is a specialized additional step of histolysis that facilitates the attachment of nascent cartilage cells to the skeletal stump. Tissue histolysis is a key event during regeneration^4,40–42^ in which tissue stiffness greatly decreases^12^. Furthermore, ECM remodeling has been hypothesized to contribute to tissue integration in regenerating newt joints^43^. It is thus likely that osteoclast-mediated tissue resorption primes the mature skeleton for tissue integration by changing the composition and rigidity of the calcified ECM so that differentiating chondrocytes can be correctly incorporated.

A second hypothesis is that skeletal resorption could help release diaphyseal periskeletal cells. These cells, together with dermal and interstitial fibroblasts, contribute to the regeneration of proximal skeletal tissues^44–47^ and are thus promising candidates to orchestrate skeletal integration. This is supported by our observations that periskeletal cells are strongly influenced by the amputation plane, as evidenced by their high and sustained expression of *RANKL* upon diaphysis amputations. The activation of *RANKL* in periskeletal cells would 1) help coordinate skeletal tissue integration in the calcified diaphysis by promoting the differentiation of myeloid immune cells into osteoclasts; and 2) stimulate osteoclast maturation directly on the surface of the calcified skeletal element.

Our work also revealed that immune responses during limb regeneration are context-dependent and adapted to promote skeletal tissue integration according to its composition. Osteoclasts derive from the myeloid lineage, in which CMPs ultimately undergo terminal differentiation into fully mature osteoclasts^8,9^. Thus, the fact that osteoclast-mediated tissue resorption is primarily triggered by diaphysis amputations demonstrates that the immune response is adapted to the injured tissues. Another adaptation of the immune system was the sustained expression of the previously uncharacterized chemokine *Loc483* in diaphysis amputations, which we demonstrated to be sufficient to promote the presence of osteoclasts in regenerating limbs. As the most closely related chemokine in humans, *CCL20*, is known to promote osteoclast differentiation^48,49^, it could be that *Loc483* may act in a similar manner in the axolotl. However, further studies are needed to elucidate how *Loc483* is regulated in diaphyseal amputations, as well as its role on osteoclastogenesis during regeneration.

Finally, our study further revealed that differences in the tissues affected by the amputation plane impact the transcriptomic profile of the AEC, such as in the expression of collagenases. This suggests that the AEC is a dynamic structure that can likewise be adapted to specific injury contexts, which agrees with our previous study showing that the AEC is important in osteoclast-mediated skeletal resorption^6^. However, it is still unclear how the AEC impacts osteoclastogenesis, and how the underlying tissue influences the AEC. One way by which an amputation-dependent AEC could influence regeneration is through the differential secretion of serine protease inhibitors, which could moderate or even terminate the histolytic process by inhibiting the conversion of MMPs into their active forms^50,51^. Intriguingly, we also observed many genes annotated as ncRNAs being differentially expressed in the AEC in our two amputation models. Given their complex biological roles^52,53^, ncRNA function in the AEC could become an exciting new field of study.

Ultimately, the fact that tissue integration mechanisms can be customized to the damaged skeletal tissues highlights both the robustness and adaptability of regeneration in the axolotl. It would thus be interesting to explore whether similar adaptations happen in other tissues that also display positional differences in cell type and ECM composition, such as the muscle^33^.

Finally, the association between the amputation position and regenerative outcomes has been reported in other models. In *Xenopus laevis*, regenerative potential depends on the tissues affected by the amputation, and regeneration efficiency correlates inversely with the ossification status of the skeletal element^54,55^. In neonatal mice, only amputations through the long bones in the limb, but not through joints, can activate chondrocyte proliferation^56^. These reports and our study thus emphasize the need to, by better understanding customized regenerative responses, start challenging the assumption that limb regeneration occurs using a one-size-fits-all molecular milieu.

## Methods

### Animal Husbandry and lines

Husbandry and experimental procedures were performed according to the Animal Ethics Committee of the State of Saxony, Germany. Animals used were selected by their size (snout to tail = ST; snout to vent = SV). All experiments in this work were performed in axolotls between 6 and 10 cm ST. Detailed animal sizes and ages for each experiment can be found in Supplementary Data 1.

Axolotl husbandry was performed in the CRTD axolotl facility using methodology adapted from^57^ and according to the European Directive 2010/63/EU, Annex III, Table 9.1. Axolotls were kept in 18-19°C water in a 12 h light/12 h dark cycle and a room temperature of 20-22°C. Animals were housed in individual tanks categorized by a water surface (WS) area and a minimum water height (MWH). Axolotls of a size up to 5 cm SV were maintained in tanks with a WS of 180 cm2 and MWH of 4.5 cm. Axolotls up to 9 cm SV were maintained in tanks with a WS of 448 cm2 and MWH of 8 cm.

White axolotls (*d/d*) and transgenic axolotl lines were used in this work. The latter included the previously published C-Ti^t*/+*^*(Sox9:Sox9-T2a-mCherry)^ETNKA^*, referred to as *Sox9-mCherry*^58^, and TgTol2(*Drer.Ctsk:eGFP*)^TSG^, referred to as *Ctsk-eGFP*^6^.

### Animal procedures

For all amputations, animals were anesthetized with 0.01% benzocaine (Sigma-Aldrich, #E1501) solution. Amputations were performed in the lower arm with a scalpel under an Olympus SZX10 stereomicroscope. Diaphysis amputations were performed in the middle of the calcified diaphysis region of the radius and ulna, whereas epiphysis amputations were conducted in the distal cartilaginous epiphysis of the same skeletal elements (Fig. 1a, b; Supplementary Fig. 1a). Staggered amputations were performed by first amputating the limbs at the level of the epiphysis. Then, a second incision using microscissors was made in the diaphysis region of one of the skeletal elements. Finally, microscissors were used to resect the tissue between the two skeletal elements, thereby separating the amputated tissue from the limb.

After surgical procedure, animals were returned to the benzocaine solution and allowed to recover for 10 min prior to being transferred back into swimming water.

In vivo skeletal staining was performed using calcein (Sigma-Aldrich, #A5533) before amputations. A 0.1% solution of calcein in swimming water was prepared and animals were submerged in this solution for 5–10 min in the dark. After staining, axolotls were transferred to a tank with clean swimming water, which was changed until the water was clear, and amputated shortly after.

Live imaging was conducted at specific time points in anesthetized animals by placing them in a 100 mm petri dish, and positioning the limb accordingly. An Olympus SZX16 stereoscope microscope (SDF Plapo 0.75xPF) with fluorescence module was used to acquire images. Length measurements of the stump skeletal elements and calcified region were performed in scaled images using the line measurement tool in FIJI.

### Tissue collection

Tissue collection was performed by euthanizing animals by immersion in a lethal dosage of anesthesia (0.1% benzocaine).

For paraffin embedding and HCR, limbs were collected and fixed in 1x MEMFa (MOPS 0.1M pH 7.4, EGTA 2mM, MgSO4 × 7 H2O 1mM, 3.7% formaldehyde) for at least 1 overnight at 4°C. For bulk RNA-seq experiments, 1 to 1.5 mm of intact diaphysis or epiphysis tissue, or tissue proximal to the amputation plane, was excised, flash frozen in liquid nitrogen, and stored at -80°C until processed for RNA extraction.

For plasma measurements, blood was collected directly from the heart immediately after euthanasia using heparin-coated pipette tips into heparin-coated tubes on ice. Then, samples were centrifuged 10 min at 3000 g, the upper phase was collected into a new heparin-coated tube, and were kept at -80°C until further processing.

### Alcian blue/alizarin red staining of 45 dpa limbs

Staining was performed as described in^7^. Images were acquired on an Olympus SZX10 stereomicroscope equipped with a color camera using cellSens Microscope Imaging Software (Evident).

### Calcium measurements in blood plasma

Calcium measurement from blood plasma were performed using a Dri-Chem NX600 (Fujifilm). 10µl of plasma from intact, diaphysis, and epiphysis amputations samples were pipetted into a Fuji DRI-Chem Slide (Fujifilm, Ca-P III #2350) and the calcium concentration was measured.

### Ctsk and Ctsk-like sequence alignments

Sequences were aligned using EMBOSS Needle Pairwise sequence alignment^59^ and processed with Sequence Manipulation Suite^60^.

### Spatial transcriptomics sequencing and analysis

Spatial transcriptomics was performed using first generation Visium Spatial Gene Expression System for Fresh Frozen tissue (10X Genomics) as previously described^61^. Briefly, *d/d* animals 8–9cm ST were amputated in diaphysis and epiphysis regions and allowed to regenerate until 3 or 5 dpa. Limbs were then harvested at the level of the upper arm, flash frozen in OCT, and then stored at -80°C. Samples were cryosectioned at -11/12 °C (chamber)/ -31°C (blade) at a thickness of 10 μm.

Optimization and gene expression assays were carried out according to the manufacturer’s instructions. Briefly, slides were fixed in −20 °C methanol, dried with isopropanol, and stained with H&E. A tile scan of all capture areas was generated using an Olympus OVK automated slide scanner system with a color camera and fluorescent module. For tissue optimization, enzymatic permeabilization was conducted for 0–30 min, followed by first-strand cDNA synthesis with fluorescent nucleotides. The slide was reimaged using the standard Cy3 filter cube. An optimal permeabilization time of 20 min was determined by visual inspection to maximize mRNA recovery while at the same time minimizing diffusion. For gene expression, the initial workflow was similar to the optimization procedure and was done according to the manufacturer’s instructions. After tissue lysis and RT, amplification of cDNA and library preparation – involving fragmentation, dA-Tailing, adapter ligation and a 18 cycles indexing PCR under following conditions: 98 °C 45 sec, 10/14 cycles [98°C 20 sec, 67°C 30 sec, 72°C 20 sec], 72°C 1 min, 4°C hold – was performed based on the manufacturer’s protocol using the Library Construction kit (10X Genomics, #PN-1000190). Included in this protocol was a double-sided SPRI bead (Beckman Coulter, #B23319) size selection (0.6x/0.8x) purification and a final purification (0.8x). After checking the quality control and quantification with Fragment Analyzer (Agilent, NGS Fragment Kit #DNF-473), the libraries were sequenced on an Illumina Novaseq6000 in paired-end mode (R1/R2: 100 cycles; I1/I2: 10 cycles), generating 50-100 million fragment pairs for each library.

The raw sequencing data was then processed with the ‘count’ command of the Space Ranger software (v3.1.1) provided by 10X Genomics. The Space Ranger reference for the axolotl genome (UKY_AmexF1_1) was built using the Merged Transcripts gene annotation file generated from the bulk RNA-seq experiments (see below, available upon request) as input for the ‘mkref’ command of Space Ranger.

For clustering and gene expression analysis, we used Seurat (v.5.2.1) and the following analysis pipeline: Barcodes (spots) are filtered by having 200 or more reads and a mitochondrial content of less than 50% and features (genes) are filtered for having at least 3 cells with at least 1 read. The raw data as UMI counts were regressed to remove the effect of library size using the log normalization. Highly variable genes were identified using the variance-stabilizing transformation (vst), selecting the top 3,000. These genes were scaled and used as input for the dimensionality reduction by PCA analysis. 50 principle components were calculated and the top 20 are used for further calculations. For visualization purposes, Uniform Manifold Approximation and Projection (UMAP, k nearest neighbors = 30) was applied to project high-dimensional gene expression data into two dimensions. Graph-based clustering (k nearest neighbors = 30, resolution = 1.0) used the Leiden algorithm^62^. Marker genes that were differentially expressed for each cluster were identified using the Wilcoxon rank-sum test and Benjamini-Hochberg method to correct for multiple comparisons (presto v1.0.0). Markers which are expressed in at least 25% of the cells in a cluster and with a Log2 fold change >2 were included and are listed in Supplementary Data 2. Data was visualized with Loupe Browser v9.0 (10x Genomics).

RNA-seq raw data (fastq) has been deposited in NCBI under the Gene Expression Omnibus (GEO) accession code GSE334433.

### Deconvolution of spatial transcriptomics

Spatial transcriptomic data were deconvoluted using cell2location (v0.1) with a single-cell reference generated from a publicly available axolotl limb scRNA-seq dataset (all samples included) previously published by Li et al.^19^ Raw sequencing data were processed using Cell Ranger (v3.1.1; 10X Genomics) with the count command and default parameters against the UKY_AmexF1_1 reference genome and the Merged Transcripts gene annotation file generated from the bulk RNA-seq experiments as before. Single-cell data were analyzed using Scanpy (v1.9.1), scvi-tools (v0.17.1), and Python (v3.9). Cells with fewer than 600 detected genes and a mitochondrial content of more than 50% were excluded. Genes detected in fewer than five cells were removed, resulting in a reference dataset comprising 25,816 genes. Apart from these filtering steps, preprocessing was performed analogously to the Visium analysis described above. Clustering was performed and the marker genes that were differentially expressed for each cluster were used as a basis to annotate each cluster (Supplementary Data 3).

Reference signatures were generated following the cell2location workflow described by Kleshchevnikov et al. ^63^ with the following parameters: genes were filtered with a cell count cutoff of 5 and a cell percentage cutoff of 0.03. A regression model was trained for 250 epochs using a batch size of 2,500 and a learning rate of 0.003. Posterior distributions were exported using 1,000 posterior samples to estimate reference cell-type expression signatures. Cell-type deconvolution in the spatial transcriptomics dataset was performed using the cell2location model with the estimated reference cell-state signatures and trained using `N_cells_per_location = 5` and `detection_alpha = 20`. Model optimization was performed for 30,000 epochs using the complete dataset and q05 output was used for representation.

### Analysis of Nfatc1 enriched spatial spots

Analysis of DEGs in *Nfatc1* expressing spatial spots was performed in Loupe Browser (v8.1.2) using the Advanced Selection tool to set up a threshold of *Nfatc1* fold expression Log2 Exp >2. This approach selected 108 barcodes/spots. These were used to run a differential expression analysis comparing the selected barcodes in the UMAP to the entire dataset. The resulting list of the top most differentially expressed gene is supplied in Supplementary Data 4.

### Analysis of Loc483^+^ population in scRNA-seq

Analysis of *Loc483*^+^ population in regenerating limbs was done by selecting all cells expressing the gene (Log2 Exp >0) in the scRNA-seq used above and applying the “Re-analyze” function in Loupe Browser (v9.0) using default parameters. Co-expression analysis was done using the “Co-expression” function in the “Features” section.

### RNA extraction, library preparation and bulk RNA sequencing

Sequencing was performed using 3 *d/d* animals (biological replicates) per amputation site and per timepoint, and each biological sample resulted from the pooling of the two amputated forelimbs of each animal. For intact samples, diaphysis and epiphysis regions were dissected from the same animal. RNA extraction was performed using RNAeasy Mini Plus Kit (Qiagen, #74134) according to the manufacturer’s instructions. Samples were disrupted and homogenized using a Polytron tissue homogenizer (Kinematica, #PT1600E) in 350 µl of RLT Plus Buffer containing β-mercaptoethanol (Sigma, #M625). Extracted RNA was stored at -80°C until processed for sequencing.

RNA sequencing libraries were prepared using Watchmaker mRNA Library Prep (Watchmaker Genomics, #7K0078) on Biomek i7 with estimated fragment sizes of 300 - 400 bp. Poly-dT pull down enrichment of mRNA was performed before sequencing 101 bp paired-end reads on an Illumina NovaSeq6000 (Illumina), generating between 50 million read pairs per sample. Bulk RNA-seq raw data (fastq) has been deposited in NCBI under the Gene Expression Omnibus (GEO) accession code GSE334432.

### Bulk RNA-seq read mapping and expression analysis

Generated reads from intact or diaphysis and epiphysis amputated limbs were mapped against the current axolotl reference genome available from NCBI (UKY_AmexF1_1; GCF_040938575.1) using HISAT2 (v2.2.1)^64^. HISAT2 was run through the command line with default parameters, and a known-splicesite-infile created from the corresponding gtf annotation file via the hisat2_extract_splice_sites.py command. StringTie (v2.2.1)^65,66^ was then run through the command line with standard parameters and the option of assembling novel transcripts to produce a Merged Transcripts annotation file that was used for transcript quantification. Finally, normalized counts per million (CPM) values for each sample were calculated using the Bioconductor package edgeR (v3.40.2) ^67^, for R (v4.2.2)^68^. Normalized gene counts (CPM) are provided in Supplementary Data 5.

### Gene Ontology (GO) analysis of differentially expressed genes

From the full list of genes found to be differentially expressed in each condition (Supplementary Data 6 and 7), ncRNAs and uncharacterized genes were filtered. The remaining genes of interest are listed in Supplementary Data 8. These were analyzed for significantly enriched GO terms via DAVID (v6.8)^69^ using default parameters, which is available in Supplementary Data 9. GO enrichment terms were considered statistically significant when *p* < 0.01.

### Paraffin sectioning and Movat’s pentachrome stainings

Histological stainings were performed in intact limbs (no amputation), and in regenerating limbs after diaphysis and epiphysis amputations at 7, and 9 dpa. 3 *d/d* animals per time point and amputation plane were used. Axolotl limbs were fixed in MEMFa and decalcified in 0.5 M EDTA for 2 weeks with daily changes of the solution. Sample embedding, sectioning and staining was performed by the CMCB Histology Facility, Dresden. Briefly, samples were dehydrated in a series of EtOH in RNase-free water until 100% EtOH, and then embedded in paraffin. Longitudinal sections of 4-5 µm were generated using a microtome. Movat’s Pentachrome (Morphisto, #12057) staining was performed according to the manufacturer’s instructions. Imaging was performed using an Olympus OVK automated slide scanner system (UPLSAPO 20x/0.75).

### Hybridization Chain Reaction (HCR) staining

Probe sets for *Ctsk*, *Ctsk-like*, *RANK*, *RANKL*, *Nfatc1*, and *Loc483* were designed using the HCR probe generator created by the Monaghan Lab (https://github.com/Monaghan-Lab/probegenerator)^70^ and purchased as oligo pools (oPools Oligo Pools) from Integrated DNA Technologies. Gene and probe sequences can be found in Supplementary Data 10. Given the similarity of their sequences, we used the entire transcript of *Ctsk* and *Ctsk-like*, including the UTRs, to design probes that were specific for each gene. We confirmed probe specificity by acquiring high-magnification images of double positive cells and verifying that the transcripts occupied different localizations in the intracellular space (Supplementary Fig. 3b, white arrowheads).

Whole mount HCR was performed according to^71^ with some modifications. Briefly, limbs from *d/d* animals were rehydrated through a series of MetOH in RNase-free water and washed three times in PBT (0.1% Tween 20 in PBS). Tissue was then delipidated in Delipidation Solution (200mM Boric acid, 4% SDS, pH 8.5 in RNAse-free water) for 2 hours at 37°C. After three washes in PBT, limbs were permeabilized with Permeabilization Solution (0.3M Glycine, 2% Triton X-100, 20% DMSO in PBS) for 1 hour at room temperature (RT). Limbs were washed again in PBT, incubated in pre-warmed Hybridization Buffer (Molecular Instruments, #BPH01726) for 5 minutes and then pre-hybridized in new Hybridization Buffer for 30 minutes at 37°C. After this, tissue was incubated overnight with Hybridization Buffer containing 2 pmol per 500 µl of probe solution. The following day, limbs were washed four times with agitation for 15 minutes with Wash Buffer (Molecular Instruments, #BPW01726) at 37°C and two times for 5 minutes in 5× SSCT (3M NaCl, 300 mM sodium citrate, 0.1% Tween 20, in water) at room temperature. Pre-amplification was performed for 5 minutes at RT in Amplification Buffer (Molecular Instruments, #BAM01826), followed by amplification for 16-24 hours at RT in Amplification buffer with 30 pmol of each hairpin. Finally, tissue was extensively washed in 5× SSCT, incubated for a minimum of one week with Hoechst 33258 (Abcam, #ab228550) 1:1000 in PBS, and cleared in EasyIndex (LifeCanvas Technologies, #EI-500-1.52) for a minimum of one overnight. Samples were mounted in a glass bottom dish in EasyIndex, and then imaged using a Zeiss LSM 980 inverted confocal laser scanning microscope (Plan-apochromat 10x/0.45) with 10 µm between optical planes using Zen acquisition software (Zeiss). Number of biological replicates and HCR repeats are described in the corresponding figure legends.

### Cloning of RNA probe for AmexColB

To create a probe for *AmexColB*, RNA was first extracted from 5 dpa regenerating limbs as previously described. cDNA was prepared using Takara PrimeScript™ 1st strand cDNA Synthesis kit (Takara Bio Inc, #6110A) according the manufacturer’s instructions and using Random 6mers primers. The probe was amplified using the primers in Supplementary Data 10 and the fragment was cloned into pGEM-T Easy Vector Systems (Promega, #A1360), according to the manufacturer’s instructions. Constructs were sequenced using the Mix2Seq Kit (Eurofins Genomics, #3094-000MSK) to select for inserts with the correct sequence. Prior to transcription, 10 μg of plasmid were linearized to obtain antisense and sense probes. For synthesizing RNA probes, in vitro transcription was carried out using T7 RNA Polymerase (Roche, #RPOLT7-RO) or SP6 RNA Polymerase (Roche, #RPOLSP6-RO), following manufacturer’s instructions. Probe sequence can be found in Supplementary Data 10.

### Whole-mount in situ hybridization (WISH)

Whole-mount in situ hybridization (WISH) was performed using in vitro transcribed digoxigenin-labelled antisense RNA probes. The protocol was adapted from^72^ and performed as in^6^. Before WISH, samples were dehydrated to 100% MetOH with serial washes of MetOH in RNAse-free water, and stored at -20°C. At the start of the protocol, samples were bleached in MetOH + 6% H2O2 at RT and fully rehydrated with decreasing concentrations of MetOH in TBST (1× TBS, 0.1% Tween 20) until 100% TBST. Tissues were then washed three times with TBST and treated with 10 μg/mL proteinase K in TBST at 37°C for 30 min. After incubation, samples were washed with TBST and rinsed with 0.1M triethanolamine (Sigma-Aldrich, #90278) in RNAse-free water (TEA) pH 7.5. Tissue was next incubated with freshly prepared 0.1M TEA + 1% acetic anhydride (Sigma-Aldrich, #320102) for 10 min and then washed again with TBST. Next, samples were re-fixed with 4% PFA + 0.2% glutaraldehyde (Sigma-Aldrich, #G6257) for 20 min and washed with TBST. TBST was removed, and samples were incubated with previously warmed hybridization solution (50% formamide, 5× SSC [3 M NaCl, 300 mM sodium citrate] (pH 5.5), 0.1% Tween 20, 50 μg/mL yeast tRNA, 100 μg/mL heparin, 1× Denhart’s, 0.1% CHAPS, 5 mM EDTA) at 65°C for 4 h. Tissue was incubated with Hybridization Solution containing the RNA probe overnight at 65°C and then washed at 65°C the following day twice with prewarmed 5× SSC (50% formamide, 5× SSC, 0.1% Tween 20), 2× SSC (50% formamide, 2× SSC, 0.1% Tween 20), and 0.2× SSC (0.2× SSC, 0.1% Tween 20) for 30 min each wash. Samples were washed with TNE buffer (10 mM Tris-HCl (pH 7.5), 500 mM NaCl, 1 mM EDTA), treated with RNase (20 μg/mL in TNE buffer) for 15 min, and washed again with TNE buffer. Next, tissue was equilibrated with MABT (100 mM Maleic acid, 150 mM NaCl, 0.1% Tween 20), blocked with MABT/Block (MABT containing 1% Blocking Reagent (Roche, #11096176001)) for 1 h at RT, and incubated with a 1:5000 dilution of alkaline phosphatase-conjugated anti-digoxigenin antibody (Roche, #11093274910) in MABT/Block overnight at 4°C. After extensive washes with MABT at RT, samples were equilibrated in NTMT (100 mM Tris-HCL (pH 9.5), 50 mM MgCl2, 100 mM NaCl, 0.1% Tween 20), and developed at RT in BM-Purple (Roche, #11442074001). Reactions were stopped with PBS and fixed with 4% PFA overnight. Probe specificity was validated by utilizing sense probe controls (negative control), which confirmed the absence of staining, alongside antisense probes (positive expression detection) that yielded a robust, localized signal. For each time point, 3 epiphysis- and 3 diaphysis-amputated limbs from different animals were processed together in the same tube and developed for the exact same amount of time. Images were acquired with an Olympus SZX10 stereomicroscope equipped with a color camera and with the cellSens Microscope Imaging Software (Evident).

### Orthology inference of Loc483

Orthology of *Loc138491483* was first examined using the aLeaves webserver^28^ with default parameters and selected databases #1 Human – Refseq, #4 Non-eutherian Mammals – Ensembl 104, #5 Non-mammalian Bony Vertebrates – Ensembl 104 and others, #6: Cartilaginous fish and cyclostomes, and #7: All vertebrate entries except mammalians in NCBI Protein.

For subsequent phylogenetic analyses, complete peptide sequences of chemokines were collected from a diverse range of jawed vertebrates, and are provided in Supplementary Data 11. Chemokines were selected based on their proximity to key marker genes (see Fig. 6g; Supplementary Fig. 7a), with the human chemokine CXCL1 included as a known outgroup to all CC chemokines.

For maximum likelihood analysis, the sequences were first aligned using MAFFT^73^ (v7.520), with the parameters: `--auto --maxiterate 10000`. The resulting alignments are provided in Supplementary Data 12. Tree construction was performed using IQ-TREE^74^ (v3.1.2), with the parameters: `-m VT+I+R4 -mtree -B 10000 -bnni -alrt 10000 --robust-phy 0.95 -wsl`. The VT+I+R4 substitution model was chosen based on the IQ-TREE implementation of ModelFinder^75^, called using the parameters: `-m MFP -mtree -B 5000 -bnni -alrt 5000 --robust-phy 0.95 -wsl`. Trimming of alignment sites was handled through the trimmed log-likelihood method^76^.

Bayesian inference trees were generated using BAli-Phy^77^ (v3.6.0), using the parameters `-S jtt+Rates.free[n=4]+inv --set infer-ambiguous-observed=true`, with 6 total chains being run. The JTT+I+R4 substitution model was used as BAli-Phy does not include the VT substitution model, and JTT+I+R4 was reported to be the second-best model by the IQ-TREE implementation of ModelFinder. BAli-Phy was run until the chains converged, which was determined using the included bp-analyze script. Phylogenetic trees were visualized using the iTOL: Interactive Tree Of Life webserver^78^ (v7).

### Injection of BAPTA and CaSO4 in regenerating limb blastemas

For the injections of 10mM BAPTA (Abcam, #ab144924), a working solution was prepared by diluting a stock solution of 25mM of BAPTA (in DMSO) in APBS (80% PBS in RNAse-free water) with 1% Fast Green (Sigma-Aldrich, #F7252) for easier visualization while injecting. For CaSO4, a 15mM CaSO4 solution in water was directly injected in the blastemas, also with 1% Fast Green for easier visualization. The concentrations of BAPTA and CaSO4 in this experiment aimed at flooding the tissue with each compound locally, thus the highest possible concentrations that did not impair regeneration were used.

Animals were first anesthetized and then injected with a Pneumatic PicoPump (WPI, #PV830) using a fine heat-pulled glass capillary to inject 150 nL of solution locally in both regenerating limbs. Ctsk^+^ signal was quantified by first defining a ROI of 3.5 mm^2^, which encompassed most of the lower arm, and applying the maximum entropy threshold method. The area of Ctsk^+^ signal was then measured using the Area function in FIJI.

### Cloning, and injection and electroporation of Loc483

To obtain and clone the coding sequence of *Loc483*, RNA was extracted from diaphysis amputated limbs and cDNA was prepared as above. The full coding sequence of *Loc483* was then amplified with Phusion^®^ High Fidelity DNA Polymerase (New England Biolabs, #M0530) from whole cDNA according to the manufacturer’s instructions and using the primers in Supplementary Data 10. The fragment contaning *Loc483* was then cloned into an vector containing the CAGGS promoter for expression using standard cloning techniques.

Plasmid administration was conducted by first anesthesizing the animals and injecting a solution of 1.5 µg/µL of *Loc483* plasmid + reporter plasmid (experimental conditions) or just the reporter plasmid (control conditions) diluted in APBS (0.8X PBS) into regenerating limbs using a fine heat-pulled glass capillary as before. Both solutions contained 1% Fast Green for easier visualization while injecting. Animals were then immediately electroporated using a Super Electroporator NEPA21 TypeII (Nepa Gene) with 2 poring pulses of 80V, 50 msec length, 50 msec interval, 9% decay rate, and 5 transfer pulses of 40V, 50 msec length, 999 msec interval, 5% decay rate, using a tweezer electrode (Nepa Gene, #CUY650P3). After electroporation, animals were returned to their holding tanks containing swimming water, and allowed to regenerate until imaged at specific time points as before. Ctsk^+^ signal was quantified by first defining a ROI of 3.5 mm^2^, applying the maximum entropy threshold method, and then measured using the Area function in FIJI.

### Image processing, analysis and quantification

All images were processed using Fiji^79^. Processing involved selecting regions of interest, merging, or splitting channels, and improving brightness and contrast levels for clear visualization in figures. Maximum intensity projections were done in all confocal images, unless otherwise specified in the figure legend. When appropriate, stitching of tiles was done directly in the acquisition software Zen (Zeiss Microscopy, Jena, Germany).

### Data representation and statistical analysis

All graphs and statistical analyses unless specified were performed with GraphPad Prism 10 (GraphPad Software, San Diego, CA, USA). Specific number of replicates, statistical tests and pos-hoc tests are indicated in the respective figure legends and in the section below. All figures and schemes were generated with Affinity Designer 2 (Serif Europe, West Bridgford, UK).

### Statistics and reproducibility

Replicates for the experiments are reported as follows. Technical replicates (total number of different animals used) are denoted as n, and independent replicates (total number of experiments performed on separate days, and/or with different batches) are denoted as N.

For HCRs per time point and amputation plane: *Ctsk/Ctsk-like*, *RANKL/Ctsk*, and *RANK/Ctsk/Nfatc1*, n = 3, N = 3; *Loc483*, n = 3, N = 4; *Ctsk/Nfatc1, Loc483/Nfatc1*, and *Ctsk/Loc483*, n = 3, N = 1. For histology analyses (Fig. 1a, 1f), n = 3 per time point and amputation plane were used, N = 2. For WISH for *AmexColB*, n = 3 per time point and amputation plane, N = 3. For resorption time course experiments (Fig 1c), n = 5 per time point and amputation plane, N = 3. For *Sox9::Ctsk* time course experiments (Fig. 2a), n = 5 per time point and amputation plane, N = 3. For calcium measurements (Fig. 3a), n = 5 per time point and amputation plane, N =4. For BAPTA and CaSO4 injections, (Fig. 3 c-h), n = 4 per time point and amputation plane, N = 2. For spatial transcriptomics (Fig. 4a), n = 2, N = 1 per amputation plane at 3 dpa. At 5 dpa, n = 2 for diaphysis, n = 1 for epiphysis, N = 1. For electroporation experiments (Fig.7), n = 3 per time point and amputation plane, N = 2. For staggered amputations experiments (Fig. 8), n =8 per time point and amputation type, N = 2. For bulk RNA-seq, n = 3 per time point and amputation plane, N = 1. All attempts at replication were successful.

## Data availability

All RNA-seq raw data generated in this study have been deposited in the NCBI Gene Expression Omnibus database under the accession code GSE334433 for spatial transcriptomics [https://www.ncbi.nlm.nih.gov/geo/query/acc.cgi?acc=GSE334433] and GSE334432 for bulk RNA-seq [https://www.ncbi.nlm.nih.gov/geo/query/acc.cgi?acc=GSE334432]. The scRNA-seq data used in this study for deconvolution and re-analysis^19^ are available in the SRA database under accession code PRJNA589484 (https://www.ncbi.nlm.nih.gov/sra/?term=PRJNA589484). No original code was generated in this work. Source Data are provided with this paper.

## Acknowledgements

We would like to thank past and current members of the Sandoval-Guzmán lab for continuous support and companionship during the development of this work. We are particularly grateful to Susanne Weiche for her excellent technical assistance, to Maximilian Krause for his valuable comments on the manuscript, and to Anja Wagner, Beate Gruhl, and Judith Konantz for their dedication to axolotl care. This work was supported by core facilities of the Technology Platform of the Center for Molecular and Cellular Bioengineering (CMCB) of the TU Dresden, namely the Genome Center, the Light Microscopy Facility, and the Histology Facility.

## Author Contributions Statement

R.A. and T.S-G. conceived the study. R.A., C.A., and T.S-G. acquired funding. R.A. designed and performed most experiments, analyzed most data, and wrote the manuscript. S.D.K., K.B., M.C., D.B.G, Y.S. and P.C. assisted with experimental work. R.A. and S.D.K. processed and analyzed bulk RNA-Seq data. R.A., U.A.F., and S.D.K., processed and analyzed spatial transcriptomic data. C.A. and A.D. advised on the project. T.S-G provided supervision, critically revised and edited the manuscript.

All authors proofread and revised the manuscript.

