## Supplementary Figures for "Tissue composition shapes differential skeletal integration strategies during axolotl limb regeneration"

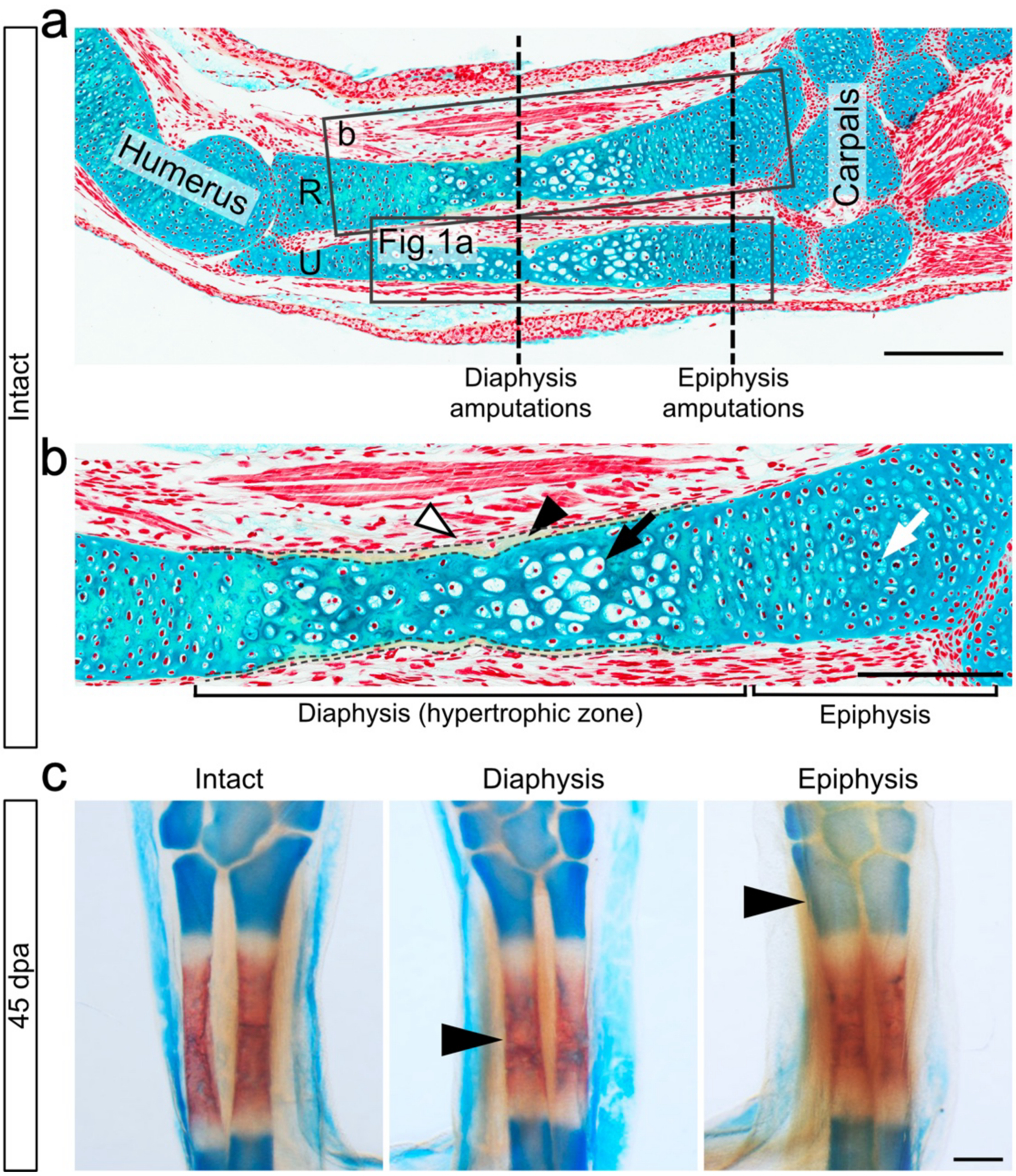


**Supplementary Fig. 1 Epiphysis and diaphysis amputations in the axolotl lower arm. a** Movat's pentachrome staining of a longitudinal section of a representative intact lower arm of an axolotl of the size ranges used in this study. Cartilaginous tissue is stained blue, calcified ECM is stained yellow, and cell bodies and nuclei are colored red. Vertical dashed lines represent the approximate site of amputation planes. Proximal is to the left, while distal is to the right of the image. Representative image of an experiment with n = 3 animals. Scale bar: 500 µm. **b** Inset of the radius in a, showing the diaphysis region, comprised by mineralized ECM (black arrowhead, in yellow between dashed lines), periskeletal cells (white arrowhead) and hypertrophic chondrocytes in the hypertrophic zone (in blue, black arrow). Epiphyseal regions are mostly composed of chondrocytes (in blue, white arrow). Scale bar: 300 µm. A similar inset of the ulna is shown in Fig.1a. **c** Representative images of Alcian Blue/Alizarin Red stainings of an intact limb (left), and limbs amputated at the level of the diaphysis (center) and epiphysis (right) at 45 dpa. The intact limb shown consists of the contralateral limb of the diaphysis amputated animal. Diaphysis and epiphysis-amputated limbs depicted here are the same as the ones shown in the time course in Fig. 2a. Black arrowheads indicate the approximate amputation site. Representative images of an experiment with n = 5 animals per time point and amputation plane. Scale bar: 500 µm.

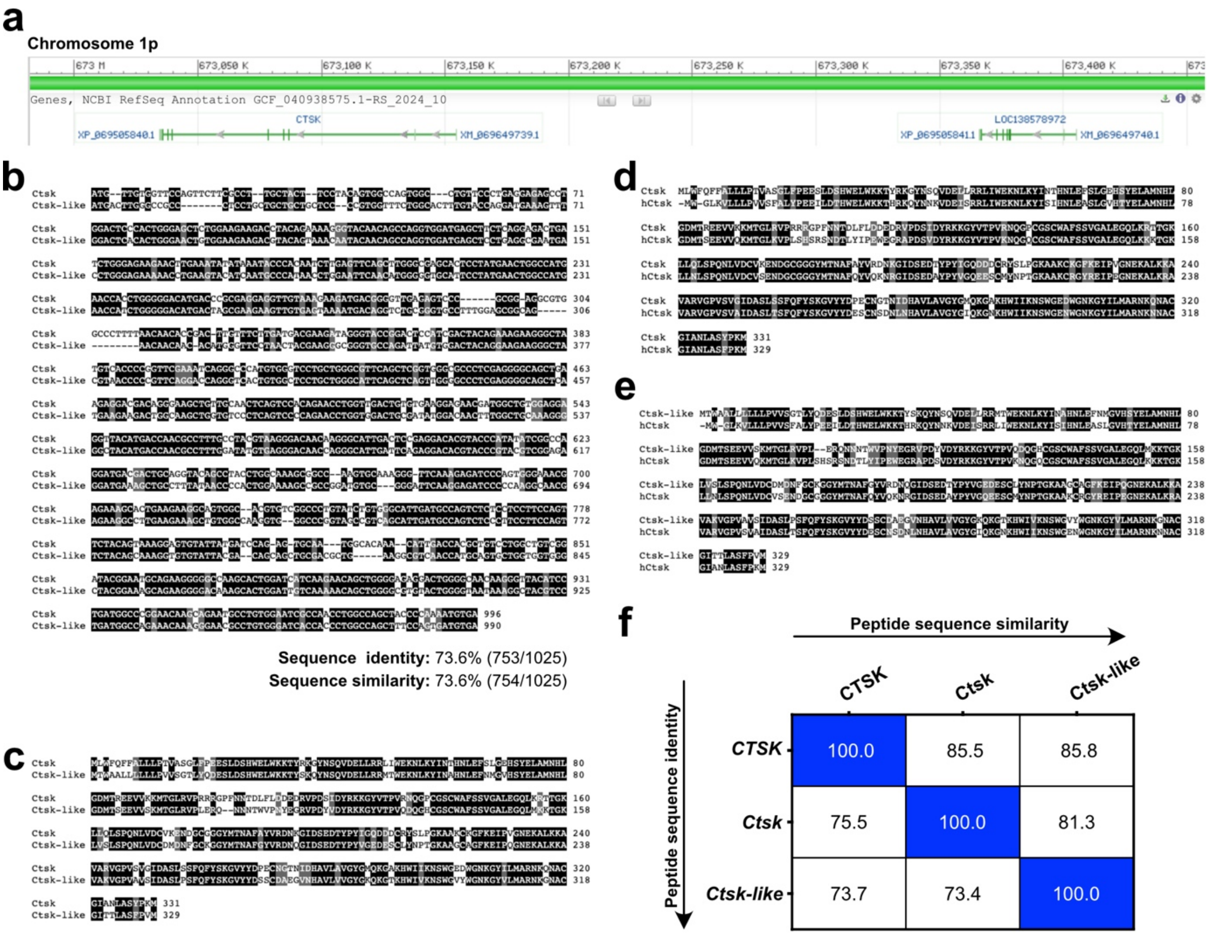


**Supplementary Fig. 2** ***Ctsk-like*** **is likely a duplication of the** ***Ctsk*** **gene present in the axolotl genome. a** Localization of *Ctsk* and *Loc138578972* (*Ctsk-like*) genes in the axolotl genome (GCF_040938575.1). **b** Alignment of predicted coding sequences of *Ctsk* and *Ctsk-like*, with sequence identity and similarity scores. **c** Predicted protein sequence alignment of *Ctsk* and *Ctsk-like*. **d** Protein sequence alignment of human *CTSK* (hCtsk) and axolotl *Ctsk*. **e** Protein sequence alignment of human *CTSK* (hCtsk) and axolotl *Ctsk-like*. **f** Table depicting calculated sequence identities and sequence similarities between human CTSK, and axolotl *Ctsk*, and *Ctsk-like* peptides.

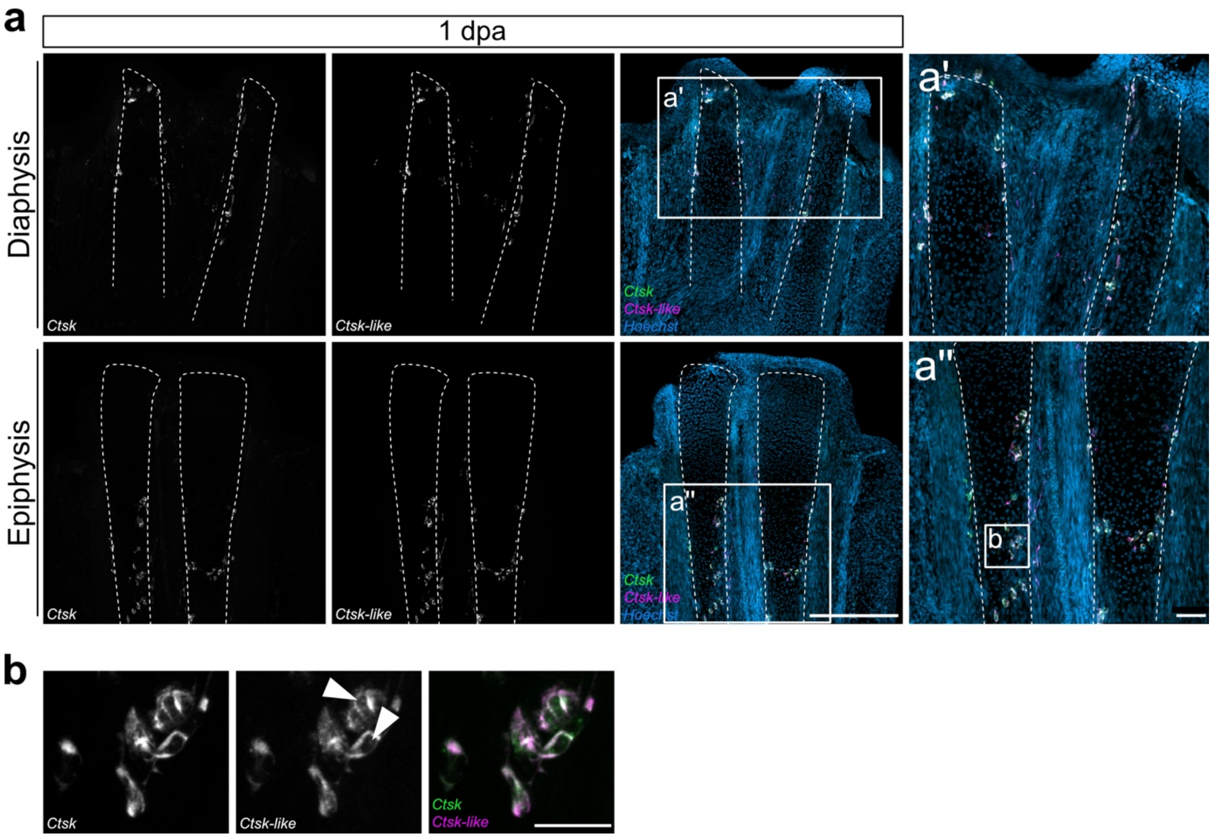


**Supplementary Fig. 3** ***Ctsk*** **and** ***Ctsk-like*** **are not differentially expressed in epiphysis and diaphysis amputations at 1 dpa. a** HCR for *Ctsk* (green) and *Ctsk-like* (magenta) at 1 dpa in representative diaphysis- and epiphysis-amputated limbs. Scale bar: 500 µm. Boxes indicate the position of the insets in a' and a''. Scale bar in insets: 100 µm. HCR was performed once in n= 3 limbs from three different animals per amputation plane. **b** High magnification of double positive *Ctsk*/*Ctsk-like* cells in a''. White arrowheads indicate intracellular spots in which the two transcripts do not overlap. Scale bar: 50 µm.

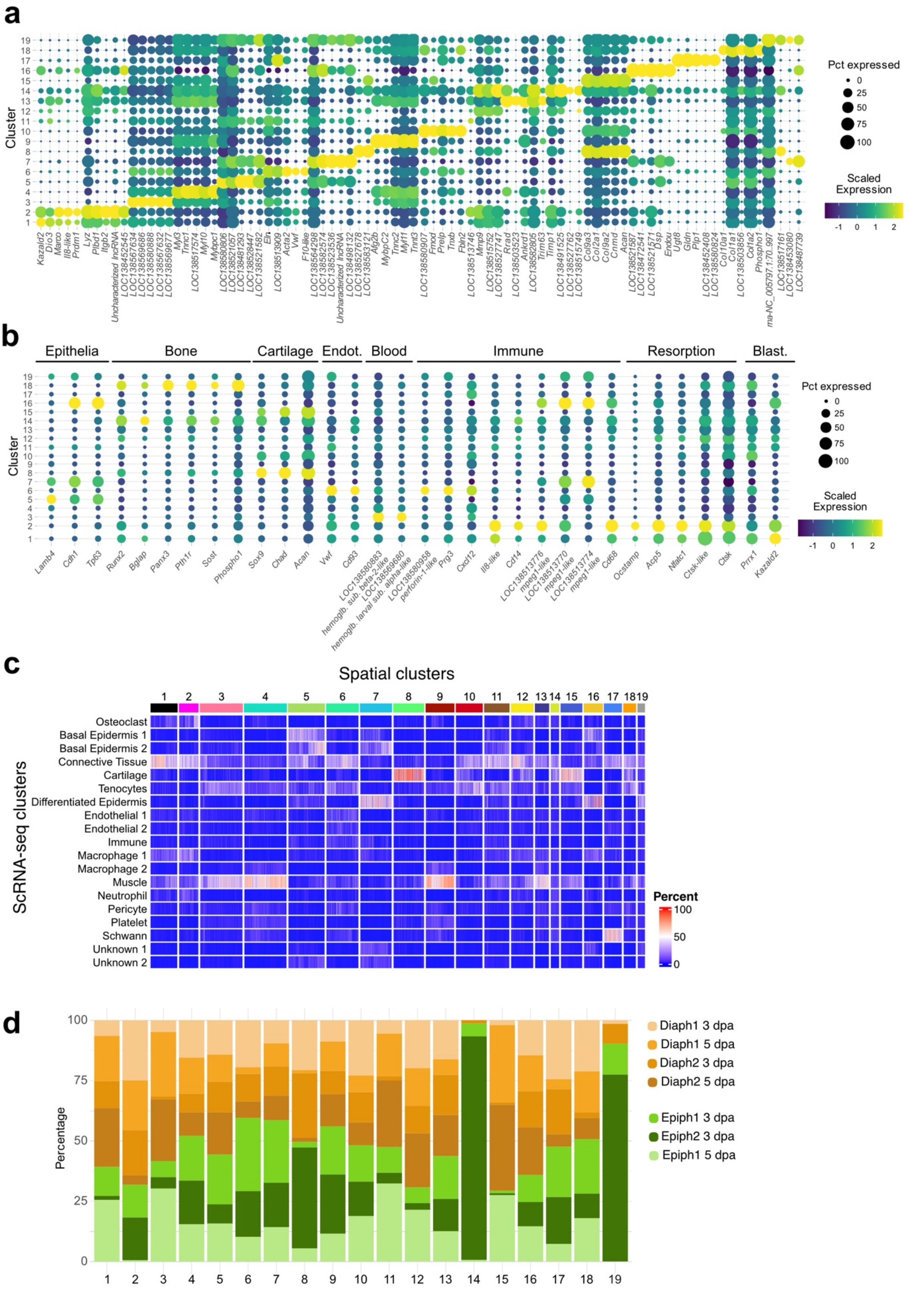


**Supplementary Fig. 4 Cluster annotation and expression of known marker genes in spatial transcriptomics. a** Dot plot showing the top marker genes in each of the 19 annotated spatial clusters. **b** Dot plot showing expression of known marker genes for epithelia, bone, cartilage, endothelium (Endot), blood, immune system, resorption and blastema (Blast) in the 19 spatial transcriptomics clusters. **c** Cell2location deconvolution of the 19 spatial clusters with cell identity annotations derived from a published scRNA-seq dataset of regenerating limbs[19]. Each vertical line represents one spatial dot, colors represent the percentage identity of each spot with the scRNA-seq categories. **d** Contribution of spatial samples to each of the 19 annotated clusters.

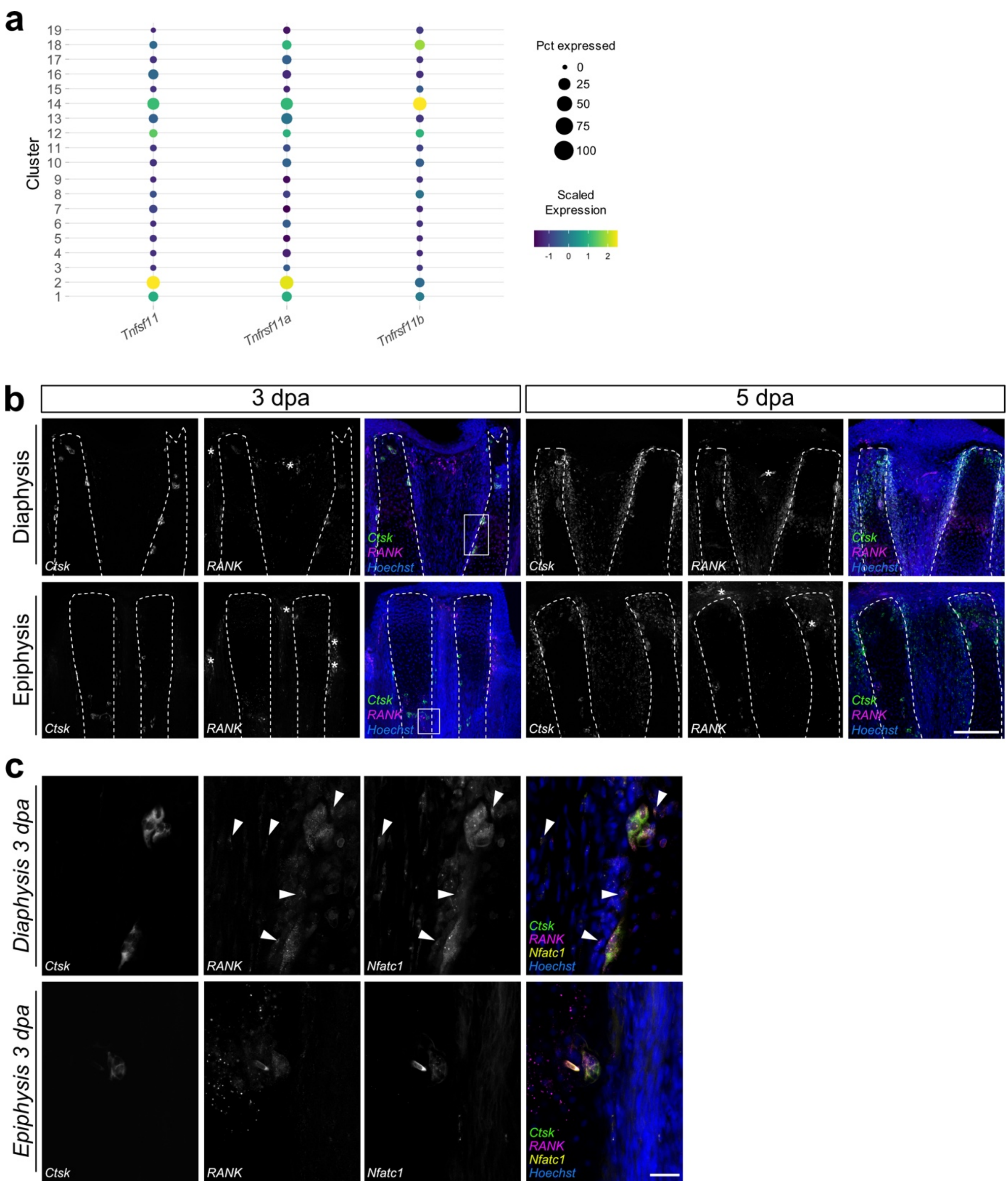


**Supplementary Fig. 5 RANK and RANKL are likely involved in osteoclastogenesis during axolotl limb regeneration. a** Dot plot showing the expression of *Tnfsf11* (*RANKL*), *Tnfrsf11a* (*RANK*), and *Tnfrsf11b* (*Osteoprotegerin*), in spatial clusters. **b** HCR for *Ctsk* (green) and *RANK* (magenta) at 3 and 5 dpa in representative diaphysis- and epiphysis-amputated limbs. Asterisks indicate autofluorescence. HCR was performed in n= 3 limbs from three different animals per amputation plane and independently repeated 3 times. Scale bar: 300 µm **c** Insets of b showing *Ctsk* (green), *RANK* (magenta), and *Nfatc1*(yellow). Arrowheads indicate cells that are positive for *RANK* and *Nfatc1*, but not *Ctsk*. Scale bar: 50 µm.

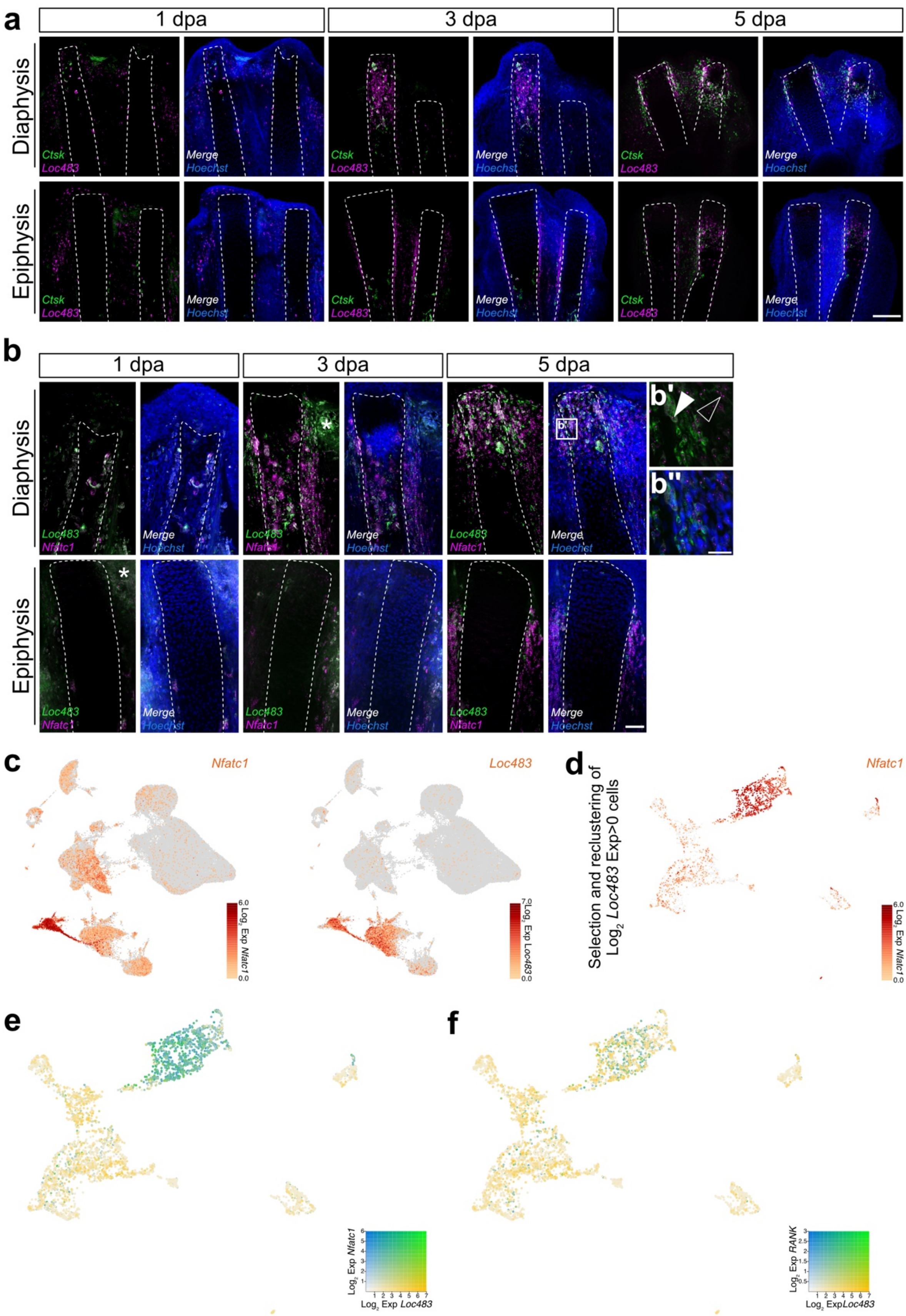


**Supplementary Fig. 6** ***Loc483*-expressing cells are a heterogenous population that can contain osteoclast progenitors. a** HCR for *Ctsk* (green) and *Loc483* (magenta) at 1, 3, and 5 dpa in representative diaphysis- and epiphysis-amputated limbs. HCR was performed once in n= 3 limbs from three different animals per time point and amputation plane. Scale bar: 300 µm. **b** HCR for *Loc483* (green) and *Nfatc1* (magenta) at 1, 3, and 5 dpa in representative diaphysis- and epiphysis-amputated limbs. Scale bar: 300 µm. Asterisk indicates background. Box indicates approximate location of insets b' and b''. Inset b' shows high magnification images of one optical section containing cells that co-express *Loc483* and *Nfatc1* (white arrowhead) or just *Nfatc1* (white-outlined arrowhead). Inset b'' shows the same as b' but with the addition of nuclear staining (Hoechst). Scale bar in insets: 100 µm. HCR was performed once in n= 3 limbs from three different animals per time point and amputation plane. **c** Expression of *Nfatc1* and *Loc483* in a UMAP of a previously published scRNA-seq dataset of regenerating axolotl limbs[19]. **d** Expression of *Nfatc1* in a UMAP created by selecting and re-clustering all *Loc483*-expressing cells from the dataset in c. Expression levels for *Nfatc1* were calculated as $Log_2$ fold expression. **e** Co-expression plots of *Loc483* and *Nfact1* in the UMAP created in d. **e** Co-expression plots of *Loc483* and *RANK* in the UMAP created in d. Expression levels for each gene were calculated as $Log_2$ fold expression.

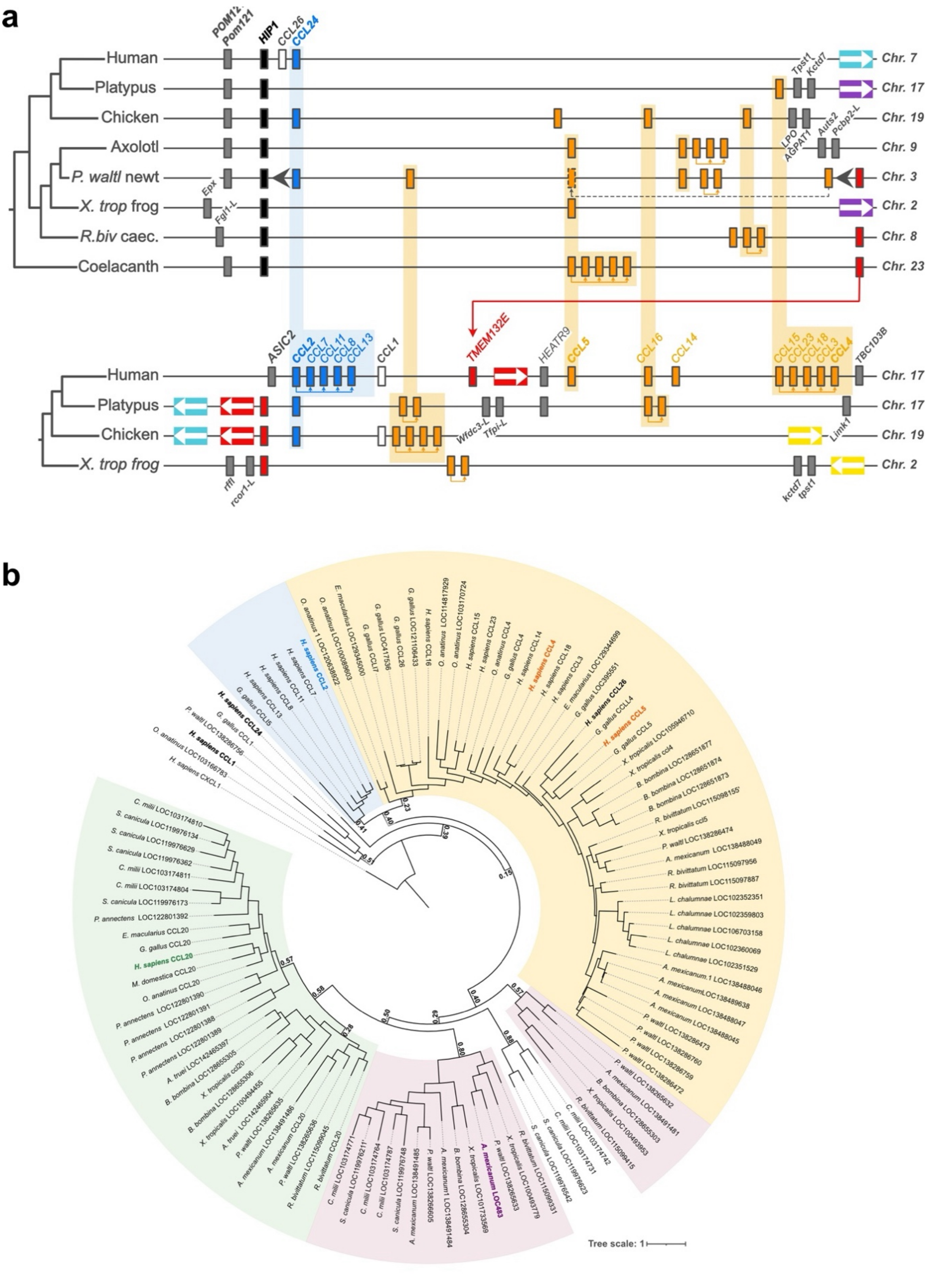


**Supplementary Fig. 7.** ***Loc483*** **is not orthologous to human** ***CCL2*****,** ***CCL4*****, or** ***CCL24*****. a** Schematic representation of genomic regions containing *Ccl2*-, *Ccl4*-, and *Ccl24*-related chemokines across several vertebrates. Non-chemokine genes *Hip1* and *Tmem132e* were used as anchor genes of the regions. Colors reflect the clades of the ML tree in Fig. 6 and the Bayesian Inference tree below. Non-chemokine genes are colored in gray, except for *Tmem132e* in red and *Hip1* in black. Orthologous genes, as determined by the ML tree, are aligned, with several chemokine clusters shaded for ease of comparison. Instances of likely tandem duplications are marked with colored arrows underneath genes. Dark grey arrowheads in the chemokine-containing region of *P. waltl* newt represent an inversion of the actual gene order for ease of orthologous gene alignment. *P. waltl* newt chemokine *Ccl5* with dashed outline is not actually present at this location, but is drawn there to clarify its orthology with other chemokines; the true location is drawn with a solid outline and connected by a dashed arrow. Colored rectangles with white arrows represent several colinear non-chemokine genes that are shared between species; colors are used to designate unique assemblages of genes, and arrows indicate their order. Gene size and distances between genes not drawn to scale. *P. waltl*, *Pleurodeles waltl*; *X. trop*, *Xenopus tropicalis*; *R. biv*, *Rhinatrema bivittatum*; caec., caecilian. **b** Bayesian Inference (BI) phylogenetic tree of chemokines selected based on syntenic similarities to axolotl *Loc483* and human *CCL2*, *CCL4*, and *CCL24*. Posterior probability support values displayed for key nodes. Clades colored based on grouping with specific chemokines: Blue – human *CCL2* & *CCL24*, Yellow – human *CCL4* & *CCL5*, Pink – axolotl *Loc483*, Green – human *CCL20*. Scale bar corresponds to the mean number of amino acid substitutions per site.

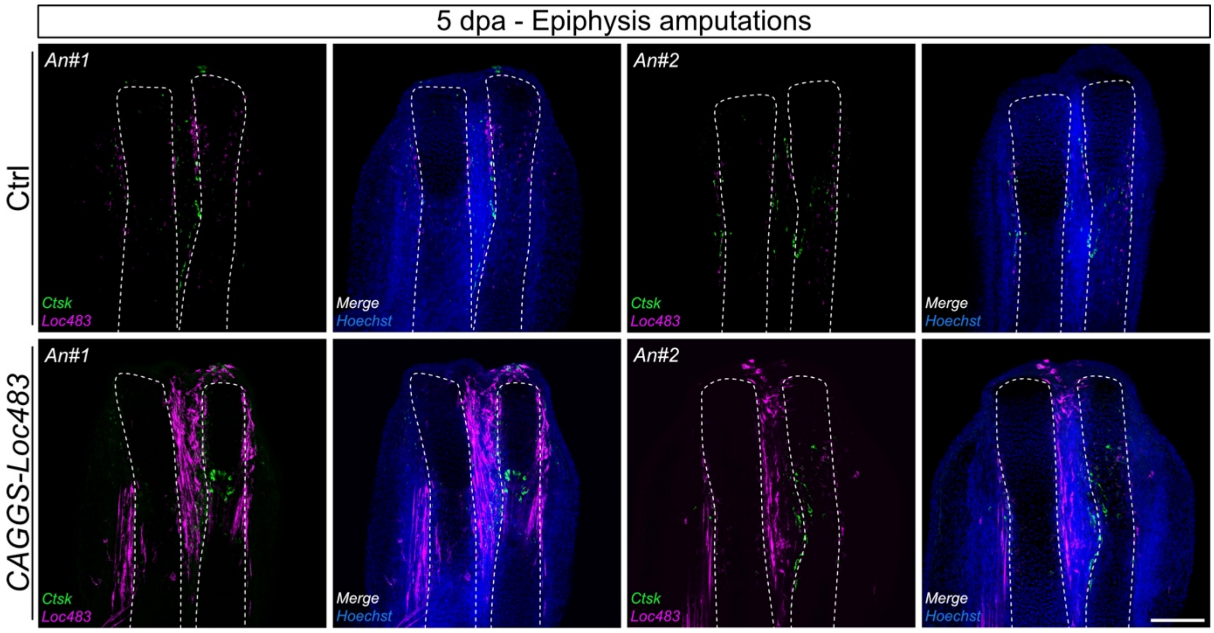


**Supplementary Fig. 8** ***CAGGS-Loc483*** **is expressed in axolotl tissues.** HCR for *Ctsk* (green) and *Loc483* (magenta) at 5 dpa in representative diaphysis- and epiphysis-amputated limbs electroporated with Ctrl or *CAGGS-Loc483* plasmid. Scale bar: 300 µm. HCR was performed once in n= 3 limbs from three different animals per amputation plane.

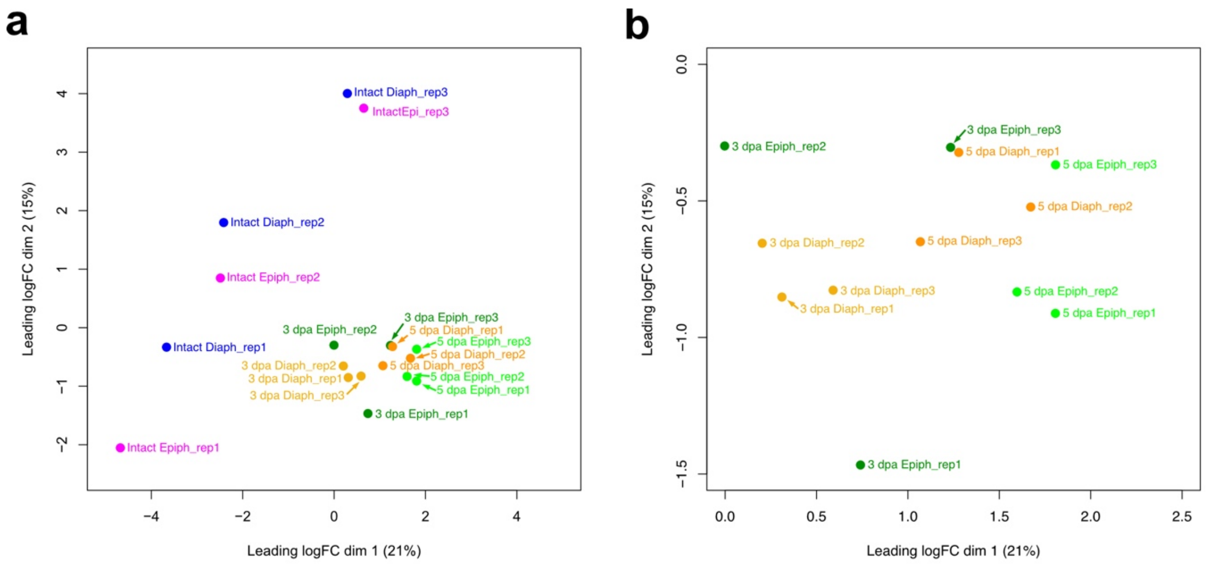


**Supplementary Fig. 9 PCA plots with the three biological replicates used per time point and amputation plane in the bulk RNA-seq. a** PCA plot containing all samples used. **b** PCA plot containing just samples of epiphysis- and diaphysis-amputated limbs at 3 and 5 dpa.

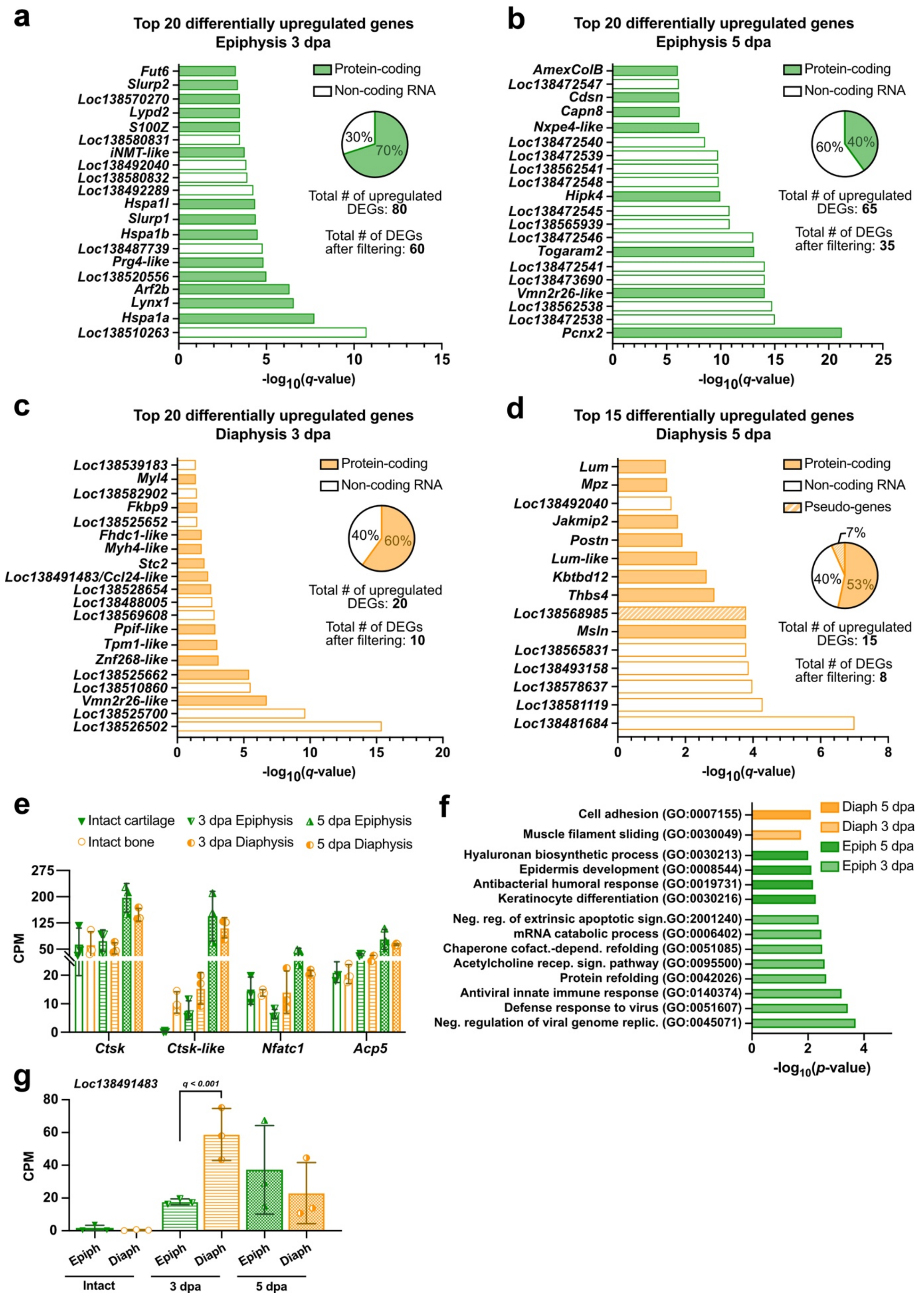


**Supplementary Fig. 10 Bulk RNA-seq in epiphysis and diaphysis amputations a** Top 20 most upregulated differentially expressed genes (DEGs), total number of upregulated DEGs, proportion of protein-coding vs non-coding DEGs, and number of DEGs after filtering used for Gene Ontology (GO) analysis in epiphysis amputations at 3 dpa. **b** Top 20 most upregulated DEGs, total number of upregulated DEGs, proportion of protein-coding vs non-coding DEGs, and number of DEGs after filtering used for GO analysis in epiphysis amputations at 5 dpa. **c** Top 20 most upregulated DEGs, total number of upregulated DEGs, proportion of protein-coding vs non-coding DEGs, and number of DEGs after filtering used for GO analysis in diaphysis amputations at 3 dpa. **d.** Top 15 most upregulated DEGs, total number of upregulated DEGs, proportion of protein-coding vs pseudo-genes vs non-coding DEGs, and number of DEGs after filtering used for GO analysis in diaphysis amputations at 5 dpa. **e** Gene expression levels of the osteoclast-associated genes *Ctsk*, *Ctsk-like*, *Nfatc1* and *Acp5* in bulk RNA-seq of intact, and diaphysis- and epiphysis-amputated limbs at 3 and 5 dpa. **f** Significantly enriched biological process GO terms for diaphysis- and epiphysis-amputated limbs at 3 and 5 dpa. $p < 0.01$, Fisher's Exact test. **g** Gene expression levels of *Loc483* in bulk RNA-seq of intact, and diaphysis- and epiphysis-amputated limbs at 3 and 5 dpa. CPM = counts per million.

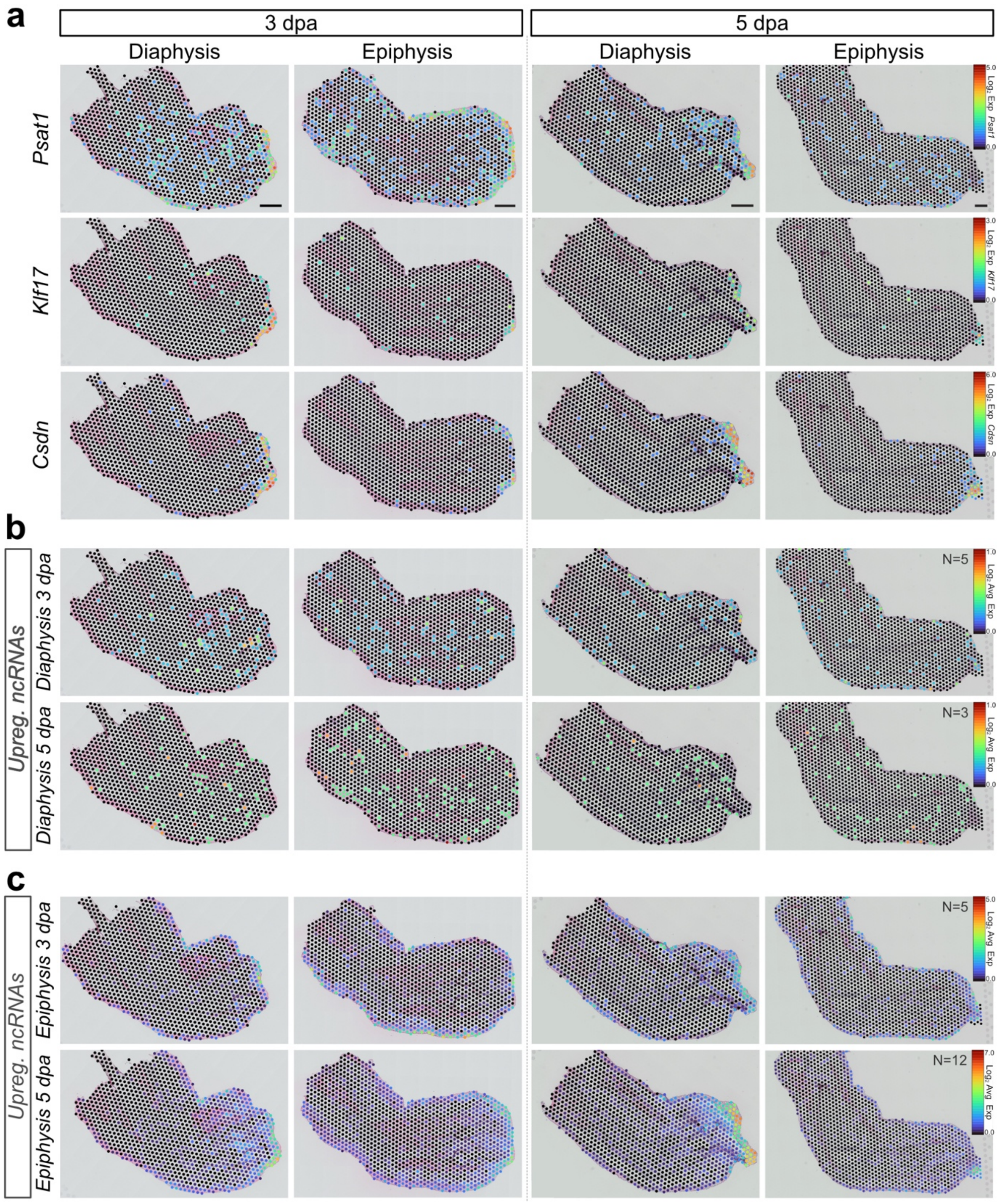


**Supplementary Fig. 11 AECs of diaphysis and epiphysis amputations exhibit differences in gene expression of non-coding RNAs.** **a** Expression of representative genes upregulated in epiphysis amputations at 3 dpa (*Psat1*) and 5 dpa (*Klf17* and *Csdn*) in bulk RNA-seq. **b** Spatial transcriptomic profile of the average expression of non-coding RNAs differentially upregulated in bulk RNA-seq of diaphysis-amputated limbs at 3 dpa (5 genes) and 5 dpa (3 genes). **c** Spatial transcriptomic profile of the average expression of non-coding RNAs differentially upregulated in bulk RNA-seq of epiphysis-amputated limbs at 3 dpa (5 genes) and 5 dpa (12 genes). Expression levels were calculated as $Log_2$ average expression. In a, expression levels were calculated as $Log_2$ fold expression; in b and c, expression levels were calculated as average $Log_2$ fold expression. Scale bar: 500 μm.

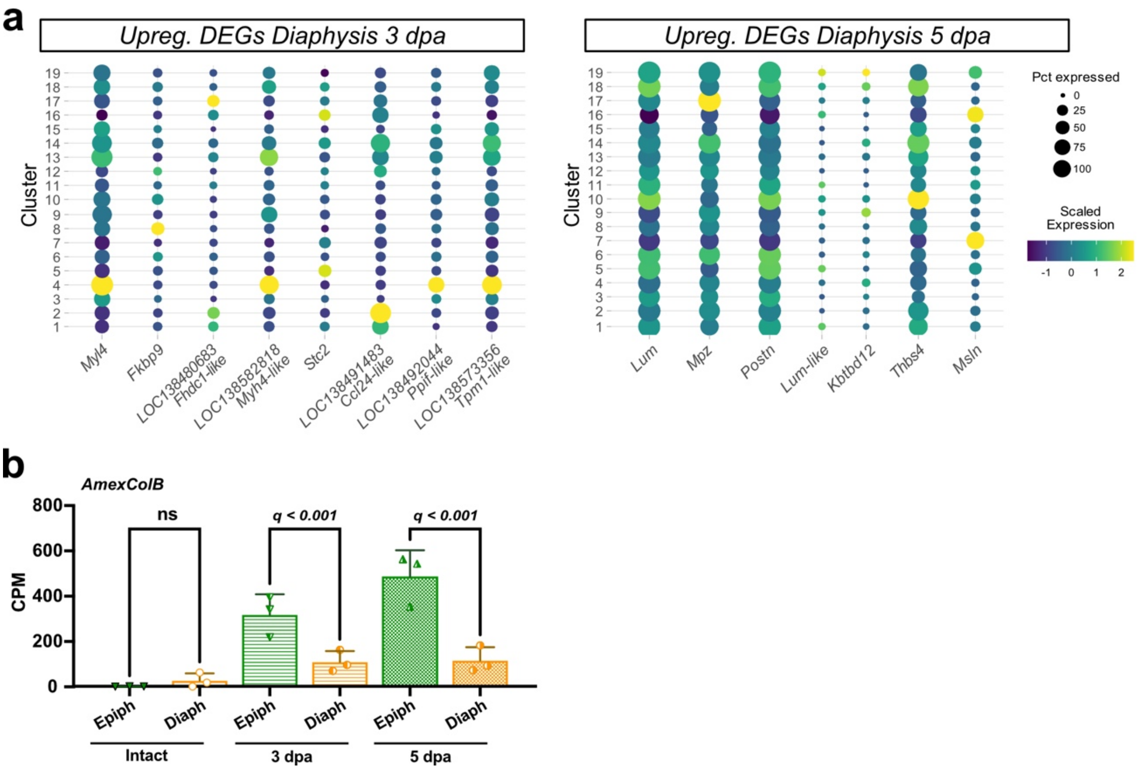


**Supplementary Fig. 12 Expression of DEGs found by bulk RNA-seq. a** Dot plots showing expression of upregulated DEGs found by bulk RNA-seq from diaphysis limbs at 3 and 5 dpa in the 19 annotated spatial transcriptomics clusters. **b** Gene expression levels of *AmexColB* in bulk RNA-seq of intact, and diaphysis- and epiphysis-amputated limbs at 3 and 5 dpa. CPM = counts per million.
